# Cellular heterogeneity in pressure and growth emerges from tissue topology and geometry

**DOI:** 10.1101/334664

**Authors:** Yuchen Long, Ibrahim Cheddadi, Vincent Mirabet, Gabriella Mosca, Mathilde Dumond, Jan Traas, Christophe Godin, Arezki Boudaoud

## Abstract

Cell-to-cell heterogeneity prevails in many biological systems, although its origin and function are often unclear. Cell hydrostatic pressure, alias turgor pressure, is essential in physiology and morphogenesis, and its spatial variations are often overlooked. Here, based on a mathematical model describing cell mechanics and water movement in a plant tissue, we predict that cell pressure anticorrelates with cell neighbour number. Using atomic force microscopy, we confirm this prediction in the Arabidopsis shoot apical meristem, a population of stem cells that generate all plant aerial organs. Pressure is predicted to correlate either positively or negatively with cellular growth rate depending on osmotic drive, cell wall extensibility, and hydraulic conductivity. The meristem exhibits one of these two regimes depending on conditions, suggesting that, in this tissue, water conductivity is non-negligible in growth control. Our results illustrate links between local topology, cell mechanical state and cell growth, with potential roles in tissue homeostasis.

**Highlights:**

- A mechano-hydraulic model of tissue growth predicts heterogeneity in cell pressure and in cell growth according to local tissue topology.
- Indentation-based measurement of cell pressure in the shoot apical meristem of *Arabidopsis thaliana*, together with a realistic mechanical model of tissue indentation, reveals pressure variations according to cell size and cell neighbour number.
- Cell growth in the Arabidopsis shoot apical meristem either correlates or anti-correlates with cell neighbour number, which can be retrieved by the model using experimentally-based parameters.

## Introduction

Cell-to-cell fluctuations are observed in many biological processes like gene expression, signalling, cell size regulation and growth [1–8]. Notably, heterogeneity in cell size and growth rate often prevails and may impact tissue patterning and macroscopic growth robustness [1,2]. Cell volume change is driven by osmosis [9–11] and the resulting intracellular hydrostatic pressure, and is restrained by peripheral constraints — plasma membrane, cytoskeletal cortex, extracellular matrix, or cell wall — in plant cells [12], animal cells [13] including tumorous [14], and microbial cells [15] (Fig. 1a).

**Figure 1.**
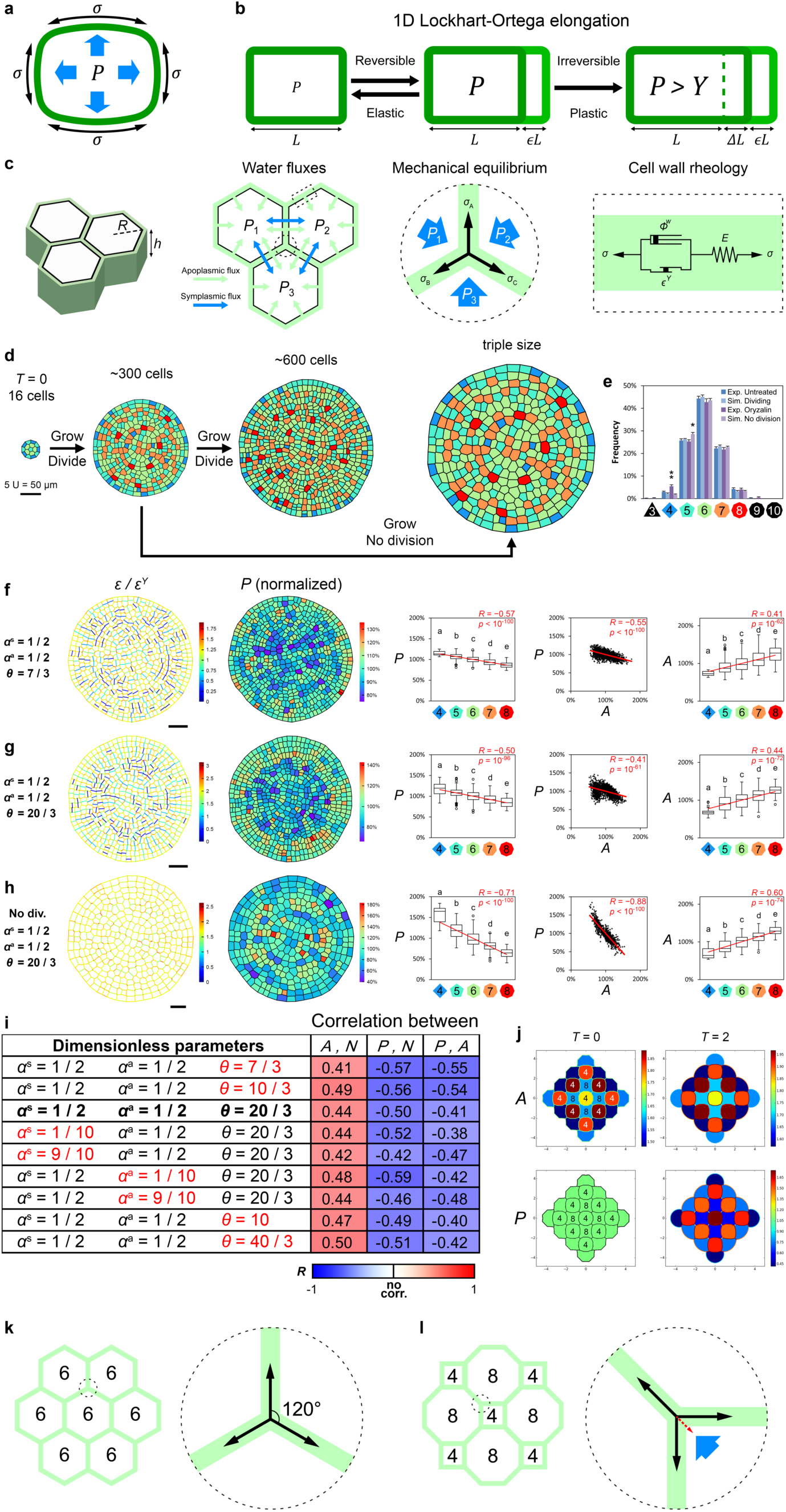
Turgor pressure heterogeneity emerges from cell topology in the mechano-hydraulic model. (**a**) In a plant cell, the turgor pressure *P* is contained by the cell wall tensile stress *σ*. (**b**) A schematic representation of the Lockhart-Ortega equation, where 1D cell length *L* elongation is a combination of reversible stretch *ϵ L* (elasticity, *ϵ* is elastic strain) and cell wall yield Δ*L* at longer timescale (viscosity) if *P* is higher than a threshold *Y* (effective plasticity, Δ*L = Ф t L* (*P* – *Y*), *Ф* is wall extensibility, *t* is time). (**c**) Schematic representations of model components, including cell geometry with height *h* and radius *R* (left), apoplasmic/transmembrane and symplasmic/intercellular water fluxes (middle left), mechanical equilibrium at tricellular junctions (middle right) and the visco-elasto-plastic cell wall rheology (right). *P*_i_, cell-specific turgor pressure; *σ*, cell wall tension; *Ф*^*w*^, wall extensibility; *ε*^*Y*^, wall strain threshold; *E*, wall Young’s modulus. (**d**) Simulation snapshots: in “dividing” simulations, 16 initial cells grow and divide until about 600 cells; in “non-dividing” simulations, divisions are stopped when cell number reaches about 300 cells and growth continues until they triple in size. Colour indicates cell neighbour number as in (e). (**e**) Similar distributions of cell neighbour number in the experimentally observed (Exp.) shoot apical meristem and in simulations (Sim.) by Kolmogorov–Smirnov test (confidence level *α* = 0.05, *D*_*n,m*_ < *D*_*α*_), error bars are standard deviations. *, Student’s *t*-test *p* < 0.05; **, *p* < 0.01. (**f-h**) Cell wall strain, turgor pressure, neighbour number, and area in dividing (f, g) and non-dividing (h) simulations; the dimensionless parameter for flux-wall balance *α*^a^, apoplasmic-symplasmic balance *α*^s^ and the osmotic drive are as indicated. Left: cell wall elastic strain *ε* normalized by yield strain *ε*^Y^; middle left: cell turgor pressure *P* normalized by average pressure; middle: boxplots of normalized cellular turgor pressure *P* against cell topology *N* (**f**, *n* = 1535 cells; **g**, *n* = 1496 cells; **h**, *n* = 759 cells); middle right: normalized pressure against normalized area; right: normalized area against neighbour number. Cells on the mesh edge were not analysed due to border effect. Circles are Tukey’s outliers; lowercase letters indicate statistically different populations (Student’s *t*-test, *p* < 0.05); red lines indicate linear regressions, with Pearson correlation coefficient *R* and corresponding *p*-value. In (**d, f-h)** scale bars are 5 unit length. (**i**) Model parameter exploration. Colours indicate Person correlation coefficient *R*, with perfect anticorrelation as blue (*R* = −1), perfect correlation in red (*R* = 1); all correlations were statistically significant (*p* < 0.05). *A*, normalized cell area; *N*, cell neighbour number; *P*, normalized turgor pressure; *α*^a^, dimensionless parameter for flux-wall balance; *α*^s^, dimensionless parameter for apoplasmic-symplasmic balance; *θ*, dimensionless osmotic drive. (**j**) Simulations initiated with 4-edged bigger than 8-edged cells, shown at two simulation times, with area and pressure shown in top and bottom, respectively; 4-edged cells have higher pressure although bigger. (**k** to **l**) Schematic explanation of topology-derived turgor pressure heterogeneity. (**k**) Three-cell junctions in a tissue of hexagonal cells are at mechanical equilibrium with equal tensions and equal wall-wall angles. (**l**) Fewer-neighboured cells have sharper wall-wall angles, a tension that is roughly constant per wall effectively results in mechanical compression due to unequal projected tension distribution (red dash-line arrow) that is balanced by higher turgor pressure build-up (big blue arrow).

Due to high difference between internal and external osmotic potential, cells with rigid cell walls – like in plants, bacteria and fungi – accumulate hydrostatic pressure, alias turgor pressure, often greater than atmospheric pressure (Fig. S1) [12]. Animal cells also accumulate hydrostatic pressure, especially when compacted or contracting [13,14], though to a lesser extent than walled cells. Whereas it is increasingly realized that pressure regulation is crucial for the general physiology, growth and signalling in animal [9,11,13] and plant cells [16–19], pressure remains poorly characterized in multicellular contexts.

In plants, turgor pressure drives cells expansion, which is classically modelled as visco-elasto-plastic process: irreversible expansion occurs when the pressure exceeds a threshold value and the cell deforms reversibly when pressure is smaller, as depicted in the Lockhart-Ortega equation (Fig. 1b) [20]:

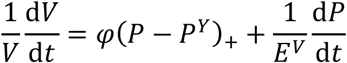

where *V* is cell volume, *P* is turgor pressure, *P*^*Y*^ is the pressure threshold for irreversible expansion, *φ* is global wall extensibility (akin to inverse of viscosity) and *E*^*V*^ is the cell volumetric modulus, while the _+_ index indicates that irreversible expansion occurs only if *P* > *P*^*Y*^. For single cell, turgor pressure positively correlates with total growth rate (Fig. 1b). Other experimental observations in single-cell systems suggest that growth rate and pressure level are not always associated [15,21], making it difficult to assess and understand the link between growth regulation and cellular pressure build-up.

In multicellular context, like the plant shoot apex and sepal epidermis, neighbouring cells grow at notably different rates [1]. This prompts the question, according to the Lockhart-Ortega equation, of whether turgor pressure also varies between neighbouring plant cells. Intuitively, pressure difference should be equalized by plasmodesmata, symplasmic bridges connecting most plant cells [22]. This is supported by the association of plasmodesmata closing (symplasmic isolation) and pressure build-up in specialized cells, such as guard cells or cotton fibres [23–25]. However, pressure gradient was also observed in plant tissues with symplasmic continuity [26], and predicted to be crucial for perception of water availability by roots [27]. In addition, computational models of tissue mechanics suggest that neighbouring cells need to have different pressure to recapitulate tissue arrangement and mechanical status in chemical-treated Arabidopsis epidermis [28] and in Drosophila epithelia [29], although such spatial variation is yet to be demonstrated, and its relation with cell-to-cell growth variability remains elusive.

Here, we explore this issue in a model plant tissue, the epidermis of the *Arabidopsis thaliana* shoot apical meristem (SAM), by combining computational modelling, dimensional analysis, and experimental observations. Our physical model that couples tissue growth mechanics with water movement accounts for our experimental observations of turgor pressure and growth rate variability in different conditions. We show that cell pressure heterogeneity emerges from local tissue topology to influence growth variability. Pressure level, however, is not a direct predictor of growth rate, and dimensional analysis of the model reveals that growth behaviour depends on the extent of the osmotic drive and on the balance between wall extensibility and hydraulic conductivity. Together, we propose a link between cell topology and cell hydro-mechanical status that may be involved in tissue homeostasis.

## Results

### A mechanical-hydraulic model predicts pressure heterogeneity emerging from tissue topology

Earlier tissue models [28,29] retrieved intracellular pressure from static tissue geometry or required differential osmotic pressure, whereas a recent model by Cheddadi et al. proposed that both pressure and growth emerge from the coupling between cell wall mechanics and classical plant hydraulics [30] in a tissue with hexagonal topology, i.e. with every cell having six neighbours. We therefore tested, based on this model, the consequences of unequal neighbour numbers on the mechanical status of the tissue, and notably on pressure (Fig. 1). We expanded on the 2D vertex model to cell walls in a multicellular tissue with more realistic 2D polygonal geometry. Generalizing the Lockhart-Ortega equation of visco-elasto-plastic 1D growth of single cell (Fig. 1b) [20], each cell wall has a thickness *w* and behaves as an elastic material of modulus *E* when wall tension induced by turgor pressure *P* is low enough, more specifically when the elastic strain *ε* (reversible deformation) is lower than a threshold *ε*^*Y*^. When tension in the wall exceeds the corresponding threshold *Eε*^*Y*^, the cell wall undergoes irreversible expansion, akin to visco-plastic flow, with an extensibility (effective viscosity) *Ф*^w^ (Fig. 1c). Water flux from extracellular space is proportional to *L*^a^Δ*Ψ*, where Δ*Ψ* = Δ*Π* – *P* is the cross-membrane water potential difference (external pressure is assumed to be zero; for parsimony we set Δ*Π* constant) and *L*^a^ is the conductivity of the cell membrane. Intercellular water redistribution via plant plasmodesmata [31], animal gap junctions, or cytoplasmic bridges [32,33] is driven by intercellular differences in turgor pressure *P*, with a conductivity *L*^s^. We did not prescribe turgor pressure, instead letting it emerge from local mechanical and hydraulic interplays (see STAR Methods for detailed model description).

To prescribe cell divisions, we utilized the recently introduced Willis-Refahi rule that derives from experimental data in the SAM: cells divide according to their size and to the size increment since the previous division [34]. The simulations recreate distributions of neighbour number (topological distributions) that are similar to those observed in the SAM (Fig. 1e).

We set the model parameters based on classical measurements from Boyer [35] and Cosgrove [36] (Table 1). We found turgor pressure to be heterogeneous, with a clear anticorrelation with topology: cells with less neighbours have relatively higher pressure (Fig 1f, 3 simulations, cell number *n* = 1535, Pearson correlation coefficient *R* = −0.57, *p* < 10^−100^).

**Table 2.** Parameters used for the untreated model. (Upper) The initial parameters correspond to *α*^s^ = 0.5, *α*^a^ = 0.5, *θ* = 7/3 in Figure 1f, which are based on measurements by Boyer (1985) [35] for *α*^a^ and *θ*, while *α*^s^ data is not available thus is conservatively ascribed an intermediate value of 0.5. (Lower) Non-dimensional parameter exploration around the reference values (initial parameters, except, *θ* = 20/3) and the corresponding modified parameters. Note that *α*^a^ = 0.9 corresponds to measurements by Cosgrove (1985) [36], compiled by Ortega (2019) [37].

| Initial parameter | Value | Initial parameter | Value |
| --- | --- | --- | --- |
| Unit length | 10 $\mu\text{m}$ | Apoplasmic | |
| Cell height $h$ | 10 $\mu\text{m}$ | conductivity $L^a$ | $3.4 \times 10^{-10} \text{ m MPa}^{-1} \text{ s}^{-1}$ |
| Cell wall thickness $w$ | 1 $\mu\text{m}$ | Symplasmic | |
| Elastic modulus $E$ | 33.8 MPa | conductivity $L^s$ | $3.4 \times 10^{-10} \text{ m MPa}^{-1} \text{ s}^{-1}$ |
| Wall extensibility $\Phi^w$ | $2.4 \times 10^{-5} \text{ MPa}^{-1} \text{ s}^{-1}$ | Osmotic pressure $\Delta\pi$ | 0.7 MPa |
| Strain threshold $\varepsilon^y$ | 0.1 | | |

| Perturbation | Modified parameter | Perturbation | Modified parameter |
| --- | --- | --- | --- |
| $\alpha^s = 0.1$ | $L^s = 3.8 \times 10^{-11} \text{ m MPa}^{-1} \text{ s}^{-1}$ | $\theta = 10 / 3$ | $\Delta\pi = 1 \text{ MPa}$ |
| $\alpha^s = 0.9$ | $L^s = 3.1 \times 10^{-9} \text{ m MPa}^{-1} \text{ s}^{-1}$ | $\theta = 20 / 3$ | $\Delta\pi = 2 \text{ MPa}$ |
| $\alpha^a = 0.1$ | $\Phi^w = 2.2 \times 10^{-4} \text{ MPa}^{-1} \text{ s}^{-1}$ | $\theta = 10$ | $\Delta\pi = 3 \text{ MPa}$ |
| $\alpha^a = 0.9$ | $\Phi^w = 2.7 \times 10^{-6} \text{ MPa}^{-1} \text{ s}^{-1}$ | $\theta = 40 / 3$ | $\Delta\pi = 4 \text{ MPa}$ |

In order to test the robustness of this result, we explored the parameter space of the vertex model. Analytical exploration in a two-cell system [30] had showed that systems dynamics is mostly controlled by three dimensionless parameters. The first, *α*^*s*^, compares intercellular water conductivity to total (intercellular and transmembrane) conductivity,

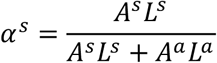

with *A*^*a*^ the average surface of a cell, *A*^*s*^ the average surface of a cell in contact with neighbouring cells, *L*^*a*^ the conductivity (per unit surface) of the plasma membrane, and *L*^*s*^ the conductivity (per unit surface) due to plasmodesmata. The second, *α*^*a*^, compares the limitation of growth by transmembrane conductivity to the combined limitation of growth by cell wall extensibility and transmembrane conductivity,

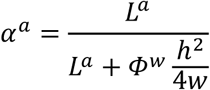

where *h* and *w* are cell height and cell wall thickness. Both *α*^*s*^ and *α*^*a*^ are bound between 0 and 1. The third parameter, *θ*, assesses the osmotic drive by comparing the cross-membrane osmotic pressure difference, Δ*Π*, and a representative threshold pressure for growth to occur, *P*^*Y*^,

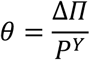

Contrary to the Lockhart-Ortega model that was formulated at cell scale, our model accounts for specific cell wall geometry and mechanical properties. Accordingly, we express the threshold in terms of a yield strain *ε*^*Y*^ and the threshold pressure depends on cell geometry and topology. We found empirically [30] that half the threshold pressure of a single hexagonal cell provides a good order of magnitude for the threshold pressure in the multicellular model and hence use

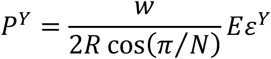

where *R* is a representative cell size and the number of cell neighbours is *N* = 6 (Fig. S1). The tissue globally grows if *θ* is greater than 1. Our first results, obtained with *α*^*s*^ = 1/2, *α*^*a*^ = 1/2, and *θ* = 7/3, corresponded to a balance between mechanical and hydraulic limitations to growth (Fig. 1f). We explored the parameter space by considering 4 values of *θ* reaching up to 40/3 (Fig. 1i, S2). As will be clarified below (see last subsection), we considered *θ* = 20/3 as a reference value (Fig. 1g). We then decreased and increased either *α*^*s*^ or *α*^*a*^ down to 0.1 or up to 0.9, respectively (Fig. 1i, S2). Note that the values 0.5 and 0.9 of *α*^*a*^ span available measurements of extensibility and conductivity [35,36] (Table 1). Next, we tested the effect of cell divisions by arresting them (Fig. 1h). In all cases, we recovered the turgor– neighbour-number anticorrelation (Fig. 1f-i, S2), demonstrating that topology-related pressure heterogeneity is a robust behaviour of the model.

In simulations, like in SAM surface, cell neighbour number and size are coupled (Fig. 1f-i, S2). Consistently, cell-specific turgor pressure anticorrelates with normalized cell area (Fig. 1f-I, S2). To uncouple cell size and topology, we modified the initial state of our simulation to have four-neighboured cells bigger than eight-neighboured cells, and found that higher turgor pressure accumulated in four-neighboured cells, not in the smaller cells (Fig. 1j), showing that turgor pressure heterogeneity emerges from tissue topology rather than cell size differences. This can be explained by local topology, which determines cell wall angles and the subsequent tension distribution at each tricellular junction (Fig. 1k, l): in our model, wall stress and strain above the growth threshold are relaxed at a rate limited by wall extensibility and hydraulic conductivity. Therefore, stress and strain are relatively homogenised in non-dividing simulations (Fig. 1h), and the sum of wall tension at each tricellular junction (vertex) mostly depends on the angles between walls. The vertex between three hexagonal cells with 120° internal angles has a sum of tension at zero (Fig. 1k). Fewer-neighboured cells have sharper internal angles, so the sum of tension at vertex is greater towards the cell interior, creating additional inward compression and prompting higher pressure build-up at equilibrium (Fig. 1l). In dividing simulations, new walls do not bear stress right after division because they form from cell interior. These new walls are gradually strained due to mesh growth but do not yield (expand) to release stress before reaching the threshold (Fig. 1f, g). Consequently, cell division keeps the wall stress from homogenizing and dampens the turgor–neighbour-number anticorrelation, compared to non-dividing simulations (Fig. 1g).

Altogether, our results imply that local hydrostatic pressure heterogeneity does not require differential cellular osmotic pressure [28] in a growing tissue, and predict a topological origin of pressure variability.

### Atomic force microscopy reveals heterogeneous turgor pressure in Arabidopsis shoot apical meristem

To test predictions, we built upon recent advances in atomic force microscopy (AFM) that enabled non-invasive turgor pressure retrieval utilizing indentation force-displacement and surface topography in living plant cells (Fig. 2) [38–42]. We used a pressurized thin-shell model to deduce the turgor pressure value from the AFM-measured force-displacement curves, which are influenced by turgor pressure, cell wall mechanical properties, cell 3D geometry [38], and may reflect mechanical properties at different sub- to supra-cellular scales according to indentation depths (Fig. 2b-e). We applied AFM measurements to the Arabidopsis SAM epidermis, a system featuring substantial growth heterogeneity [1]. We included untreated soil-grown SAMs (Fig. 3) and a conceptually simpler model SAM co-treated with naphthylphthalamic acid (NPA), a polar auxin transport inhibitor that induces pin-formed SAMs, and oryzalin, a microtubule-depolymerizing drug that blocks cell division but permits continuous, isotropic growth; hereafter referred to as “oryzalin-treated SAMs” (Fig. 4) [28]. Walls between cells are often curved in growing oryzalin-treated SAMs, suggesting that neighbouring cells have different turgor pressure [28] in a growing tissue.

**Figure 2.**
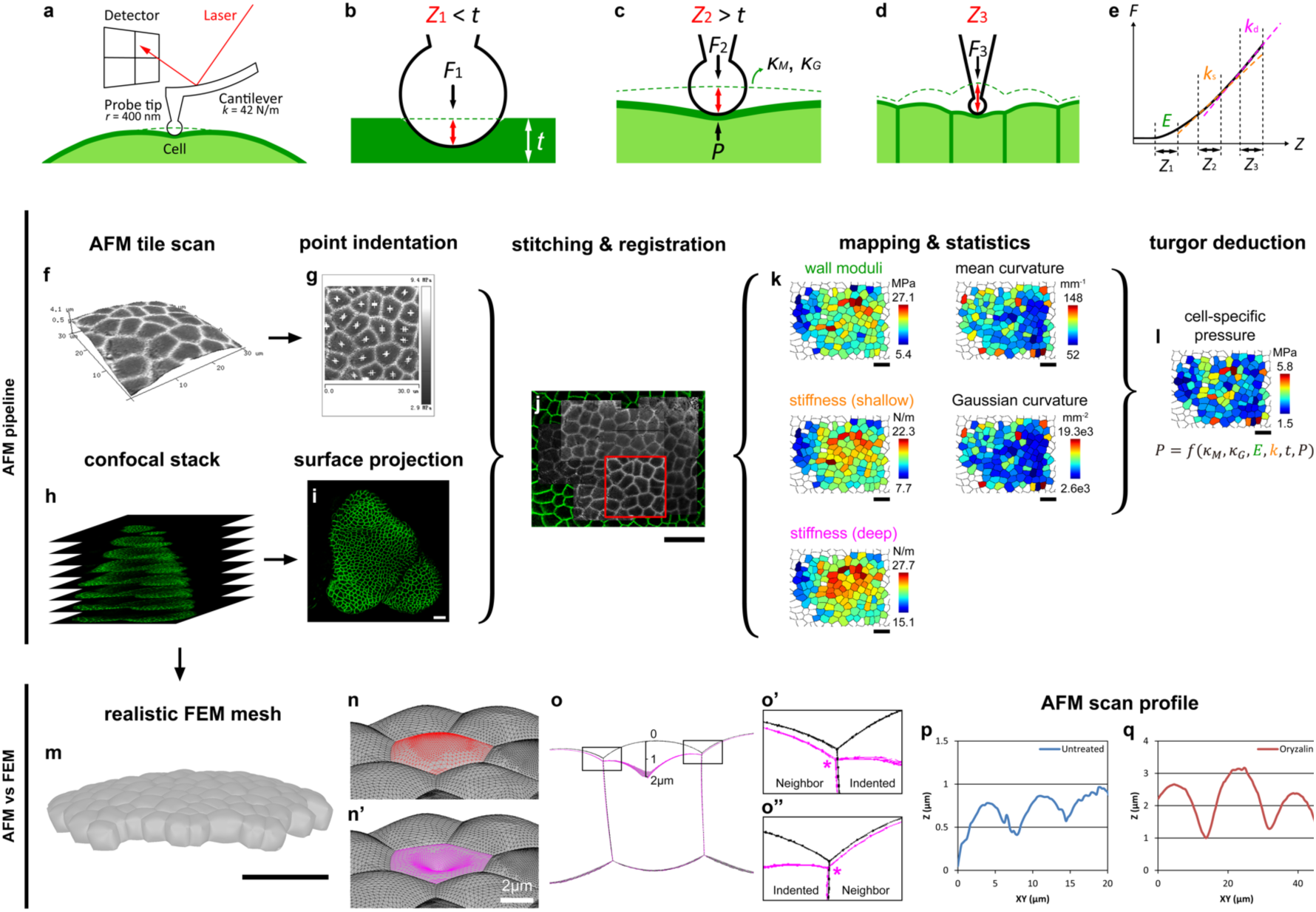
The experimental pipeline for turgor pressure deduction. (**a**) Schematic representation of AFM nanoindentation for turgor pressure measurement. *r*, probe tip radius; *k*, cantilever stiffness. (**b** to **e**) Illustration for force curve interpretation at different indentation depth. (b) When indentation depth *Z*_1_ is smaller than wall thickness *t*, the force-indentation curve is sensitive to cell wall property, (c) deeper-than-wall indentation *Z*_2_ > *t* is also sensitive to turgor pressure *P*. (d) Even deeper indentation *Z*_3_ deforms surrounding cells and is also sensitive to tissue context. Dotted line marks the shell position before indentation, which is used to determine surface mean and Gaussian curvature, *κ*_M_ and *κ*_G_, by AFM topographic scan. (e) Three regimes of the force-indentation curve are used to fit for cell wall Young’s modulus *E*, apparent stiffness at cell-scale *k*_s_ and tissue scale *k*_d_. *F* denotes indentation force. (**f** to **l**) The AFM-confocal pipeline of measurement and deduction on an example untreated SAM. (f) Derjaguin-Muller-Toporov (DMT) modulus map, highlighting cell contours, is projected on the surface topography of an AFM scan. Smaller scan region is chosen to bypass the SAM surface unevenness. (g) Multiple indentations (marked by crosshair) are performed near the barycentre of each cell. (h and i) Confocal stack and its surface projection of the same SAM with plasma membrane GFP signal. (j) Tiled AFM scans are overlaid and stitched on the confocal surface projection image, red square marking the same tile from (f and g). Individual indentation positions are registered on a global coordinate and assigned to segmented cells. (k) Force curves are analysed, and the cellular average of physical values are mapped. (l) Turgor pressure is deduced per force curve and averaged per cell. Stiffness from very deep indentation is not used for cell-specific deduction. (**m** to **o”**) Virtual indentation on realistic 3D mesh. (m) The epidermal layer is meshed based on confocal image. Cells are pressurized with a uniform 2 MPa turgor pressure. (n and n’) A cell before (red) and being indented (magenta). (o to o”) Longitudinal section of the indented mesh; black, before indentation; magenta, being indented. Cell junctions (rectangles) are magnified to highlight the neighbour cell deformation (marked by asterisks) by very deep indentation. (**p** and **q**) AFM-determined surface profiles of untreated (p) and oryzalin-treated SAM (q). Scale bars represent 20 µm if not specified. Note that z axes in (f, p and q) are stretched to emphasize SAM surface unevenness.

**Figure 3.**
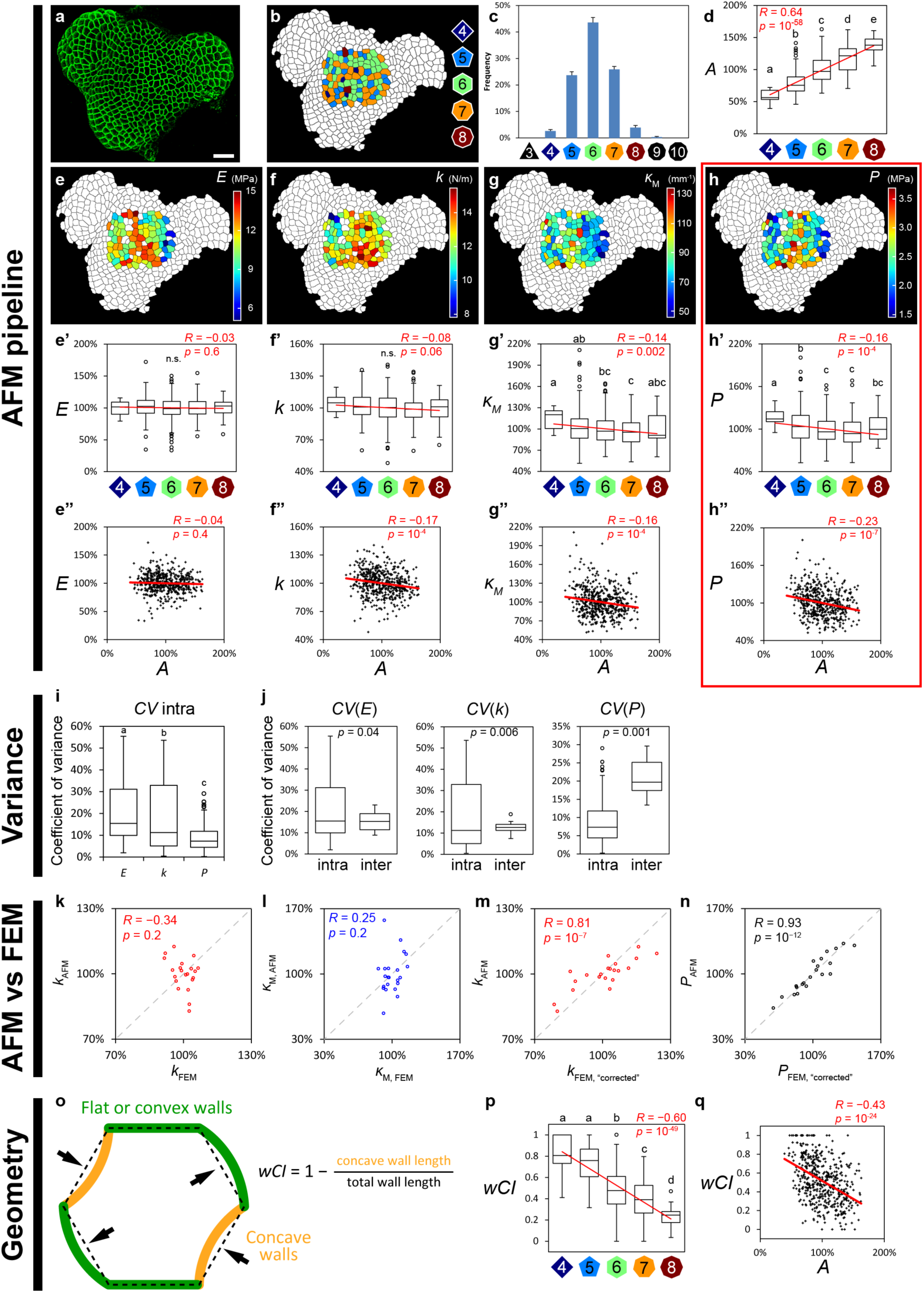
Turgor pressure heterogeneity in untreated SAM. (**a**) Top-view surface projections of untreated SAM with plasma membrane GFP signal; scale bars represent 20 µm. (**b**) Map of epidermal cell neighbour number. (**c**) Distribution of neighbour number on 7 untreated SAM surfaces, error bars are standard errors of means. (**d**) Normalized cell area *A* and cell neighbour number *N* are linearly correlated. (**e** to **h”**) AFM-determined cellular values and their association with neighbour number *N* and cell area *A*, all plotted values are normalized per SAM (7 SAMs, *n* = 503 cells). (e to e”) Cell wall Young’s moduli *E*, (f to f”) apparent stiffness *k* at intermediate indentation depth, (g to g”) cell surface mean curvature *κ*_M_, (h to h”) deduced turgor pressure *P*. (**i** and **j**) Coefficient of variance (CV) of AFM-determined values in untreated meristems. (i) At subcellular scale (intra), *P* is much less variable than *E* and *k*. (j) *P*, unlike *E* and *k*, are more heterogeneous between cells (inter) than between measurements within single cells (intra). (**o** to **q**) Weighed convex index (wCI) anticorrelates with cell neighbour number (p) and normalized cell area (q). In box plots, circles are Tukey’s outliers; lowercase letters indicate statistically different populations (Student’s *t*-test, *p* < 0.05); red lines indicate linear regressions, with Pearson correlation coefficient *R* and corresponding *p*-value.

**Figure 4.**
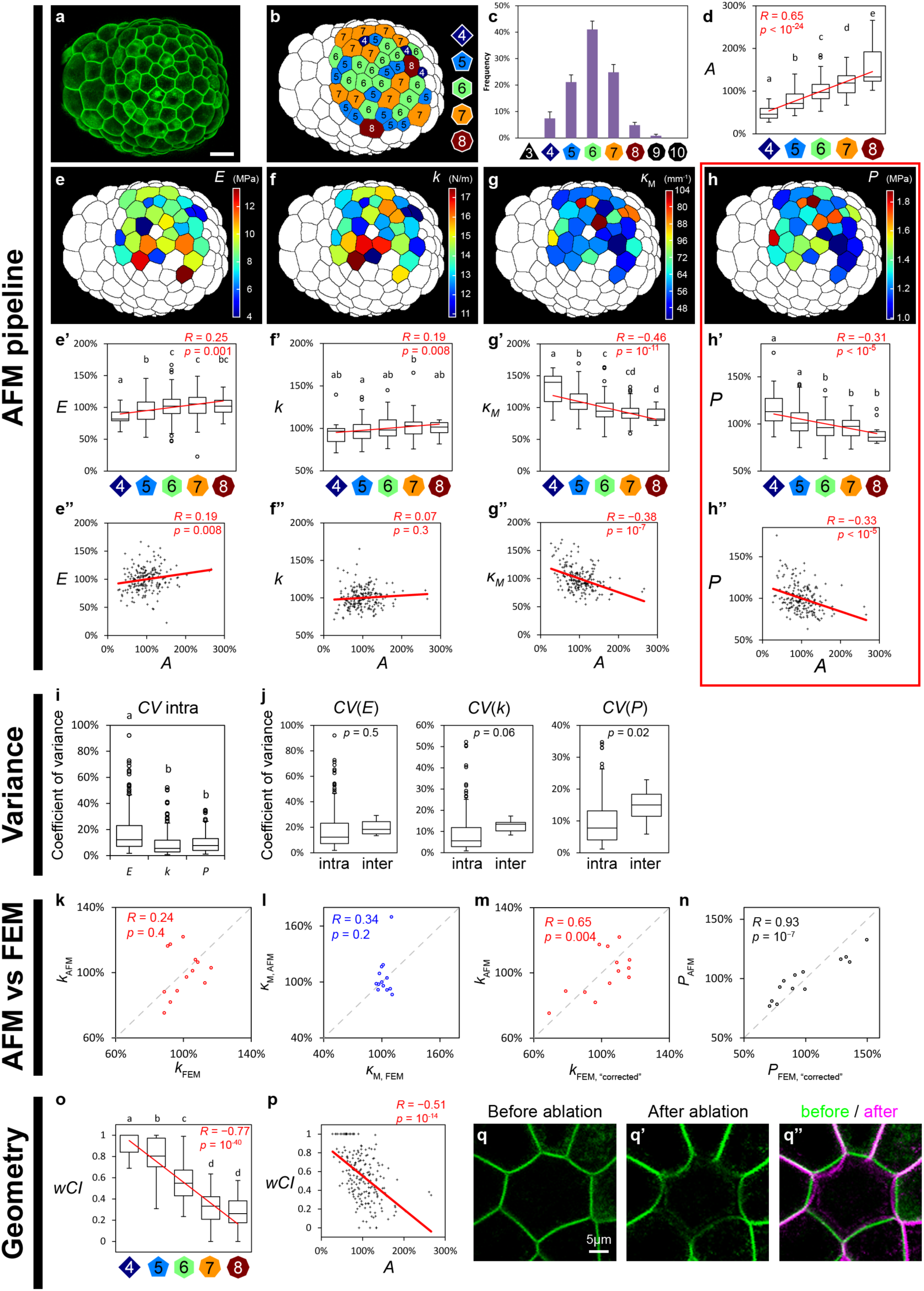
Turgor pressure heterogeneity in oryzalin-treated SAM. (**c**) Top-view surface projections of oryzalin SAM with plasma membrane GFP signal; scale bars represent 20 µm. (**b**) Map of epidermal cell neighbour number. (**c**) Distribution of neighbour number on 9 untreated SAM surfaces, error bars are standard errors of means. (**d**) Normalized cell area *A* and cell neighbour number *N* are linearly correlated. (**e** to **h”**) AFM-determined cellular values and their association with neighbour number *N* and cell area *A*, all plotted values are normalized per SAM (9 SAMs, *n* = 202 cells). (e to e”) Cell wall Young’s moduli *E*, (f to f”) apparent stiffness *k* at intermediate indentation depth, (g to g”) cell surface mean curvature *κ*_M_, (h to h”) deduced turgor pressure *P*. (**i** and **j**) Coefficient of variance (CV) of AFM-determined values in untreated meristems. (i) At subcellular scale (intra), *P* and *k* are much less variable than *E*. (j) *P* and *k*, unlike *E*, are more heterogeneous between cells (inter) than between measurements within single cells (intra). (**o** to **q”**) Weighed convex index (wCI) anticorrelates with cell neighbour number (o) and normalized cell area (p). (q to q”) Cell walls curve towards cell with low pressure, as signified by the enhanced curvature towards the ablated cell with released pressure. In box plots, circles are Tukey’s outliers; lowercase letters indicate statistically different populations (Student’s *t*-test, *p* < 0.05); red lines indicate linear regressions, with Pearson correlation coefficient *R* and corresponding *p*-value.

Figure 2 illustrates the AFM measurement pipeline, where we determined SAM surface topography with AFM (Fig. 2f) and performed indentations near the cell centre to have near-perpendicular indentation and minimize any bias due to surface slope (Fig 2g). Physical values required to deduce pressure using the published formula [39,40], except cell wall thickness separately measured by electron microscopy (EM), are simultaneously determined by AFM scan and indentation. Specifically, both mean curvature *κ*_M_ and Gaussian curvature *κ*_G_ of outer cell walls directly contribute to the capacity to sustain turgor pressure [38,39,43], and were determined from AFM scan topography (Fig. 2f). As previously suggested in tomato SAM [44], outer periclinal wall thickness is not very variable between cells or within cells in Arabidopsis SAM, with untreated meristem *t*_u_ = 179 ± 7 nm (mean ± standard error of mean, SEM) and oryzalin *t*_o_ = 742 ± 29 nm (Fig. S3). Based on previous work [45], we used the indentation depth range 0 to 150 nm to determine apparent Young’s modulus of cell wall, *E* (Fig. 2b, e). We determined indentation stiffness, *k*, using the depth range 0.3 to 0.5 µm for untreated and 1.1 to 1.5 µm for oryzalin-treated meristems (Fig. 2c, e); these ranges were chosen so that depth is greater than wall thickness (to minimize cell wall contributions to *k*) and the deformed region remains smaller than cell size (to minimize the contributions of neighbouring cells to *k*, Fig. 2d, e), so as to be in the validity range of the pressurized shell model [40]. Note that these values of depth for untreated SAMs differ from the larger values (1 to 2 µm) used in preliminary experiments [41], which were interpreted using supracellular curvature (unlike here) and were chosen to reveal supracellular pressure (averaged over many cells and possibly over cell layers) for comparison with the large-scale pressure obtained using indentation with a large flat tip (100 µm diameter). To verify the validity of the depth ranges used here, we tested influence of tissue context on indentation readout [46] by numerical simulations of indentations on realistic 3D meshes accounting for pressurized epidermal cells (Fig. 2m-o”); in conditions corresponding to experiments, the deformation of neighbouring cells was negligible, demonstrating that the depth range enables cell-level measurements.

Cell-specific AFM indentations on seven untreated and nine oryzalin-treated SAMs revealed that surface wall curvature, Young’s moduli, cell apparent stiffness and the deduced turgor pressure are all markedly heterogeneous across SAM epidermis (Fig. 3, 4). We computed intracellular variability of quantities measured at subcellular scale (which excludes curvature, measured at cell scale), and found that apparent Young’s modulus *E* and stiffness *k* show significantly higher variability than pressure *P* (as quantified by intracellular coefficients of variation, Fig. 3i, 4i), which agrees with previous work [40], and supports deduced *P* being a cellular property. Additionally, we found that *E* shows relatively comparable subcellular and intercellular variability, as previously observed [45,47], whereas *P* show significantly bigger heterogeneity between than within cells (Fig. 3j, 4j). Deduced pressure, *P*, is relatively insensitive to variability in thickness as described in [40], and change in thickness by 1× standard deviation (SD) only alters the coefficient of variance (CV) of *P* by 3% and 9% for untreated and oryzalin-treated samples respectfully, both significantly smaller than the intercellular *P* variability (untreated 21% and oryzalin 15%). All these indicate that *P* deduction is cell-specific, and that variability in cell wall mechanics does not account for deduced pressure heterogeneity.

### Osmotic treatments reveal large values of osmotic pressure in the shoot apical meristem

Based on AFM, we find that deduced turgor pressure is heterogeneous, with averages values per meristem of 2.61 ± 0.03 MPa and 1.21 ± 0.34 MPa (mean ± SEM) in untreated and oryzalin-treated meristems, respectively (Fig. S3). These values appear high with respect to the classical 0.3 to 1 MPa range found in most other plant tissues, though values of up to 5 MPa were measured in guard cells of other species using the pressure probe [48].

As neither the classic pressure probe nor the pico gauge [49] can be applied to cells as small as in the shoot apical meristem, we assessed quantitatively the values of turgor pressure *P*, by using an incipient plasmolysis assay to determine the osmotic pressure of SAM, and found that untreated meristems had a rather invariable osmolarity of about 0.5 Osm, slightly higher than values reported in tomato SAM [50], while oryzalin samples showed a wider variability of 0.6 to 1.0 Osm (Fig. S4). The corresponding values of osmotic pressure, 1.2 MPa for untreated and 2.0 MPa for oryzalin treated, are comparable to the values of AFM-deduced turgor (2.6 MPa and 1.2 MPa, respectively) (Fig. S3, S4). The quantitative agreement is less good in the case of untreated meristems, possibly due to tissue tension in the epidermis which could artificially increase the deduced pressure due to contributions from inner layers. We note however that tissue tension would smooth out intercellular differences and would thus not contribute to the cell-to-cell heterogeneity of pressure. An alternative explanation would be that solute penetration differs between untreated and treated SAMs. Altogether, the values of turgor found with AFM are in semi-quantitative agreement with the values of osmotic pressure deduced from incipient plasmolysis.

### Cell pressure in shoot apical meristem anticorrelates with local topology and size

Next, we tested model predictions and found that, indeed, AFM-determined cellular pressure anticorrelates with the number of epidermal cell-neighbours *N*, or local topology (7 SAMs, *n* = 503 cells, *R* = −0.16, *p* = 10^−4^, Fig. S5). With the linear relationship between cell area and neighbour number (Fig. 3d) due to fundamental geometrical constraints in compact tissues [51], we also identified strong anticorrelation between pressure and cell size *A* (*R* = −0.19, *p* = 10^−5^). We focused on cell-to-cell turgor variations and we removed variations between SAMs by normalizing cellular pressure to the average pressure per SAM; given that untreated SAMs have unvaried average turgor values (Fig. S3), the anticorrelation is conserved for pressure-versus-topology (*R* = −0.16, *p* = 10^−4^) and versus-size (*R* = −0.23, *p* = 10^−7^) (Fig. 3h-h”).

Encased in rigid cell walls, plant cells seldom exchange neighbours, and the main source of topological change is via division, where dividing cells tend to lose neighbours and cells adjacent to the division plain tend to gain neighbours [52]. We therefore considered oryzalin-treated SAM, in which cell divisions are arrested while growth is continuous, and recovered similar anticorrelation for pressure against neighbour number (9 SAMs, *n* = 202 cells, *R* = −0.19, *p* = 0.008) and against cell area (*R* = −0.48, *p* = 10^−12^) (Fig. S5). Normalization further enhanced the pressure-versus-topology anticorrelation (*R* = −0.31, *p* < 10^−5^) while preserving the pressure-size trend (*R* = −0.33, *p* < 10^−5^) (Fig. 4h-h”), indicating that the pressure-topology and pressure-size relationships are robust system-level trends, regardless of the absolute values.

We then examined whether such trends may be caused by trends in cell wall thickness, *t*, or modulus, *E*, or by trends in stiffness, *k*. There is no correlation between *t* and cell size (*R* = −0.10, *p* = 0.65, Fig. S3). So *t* only introduces small unbiased noise, and it cannot account for the *P* heterogeneity. We found that *E* and *k* showed no significant correlation to cell neighbour number *N* in untreated SAMs (Fig. 3e-f’), indicating that they introduce no systemic bias to *P* heterogeneity. In oryzalin-treated SAMs, however, both *E* and *k* showed weak positive correlation to *N* (Fig. 4e-f’), which is opposite to the *P* vs *N* anticorrelation (Fig. 4h’). This is interesting, because the higher *P* in *N* = 4 cells cannot be explained by lower *E* and *k* (“softer” cell). This indicates that, although feedback from shape on *E* may occur in oryzalin-treated meristems, *P* heterogeneity is not a direct consequence of feedbacks from wall tension or cell geometry/topology on wall stiffness.

Moreover, we found that turgor pressure heterogeneity may be removed when sample is osmotically challenged: the same SAM shows heterogeneous pressure when turgid and homogeneous pressure when at intermediate turgidity (Fig. S4). This shows that the AFM approach is not technically biased by tissue topology and/or cell size.

Finally, we used cell side wall convexity as a proxy for differences in turgor, because cells with higher pressure would be expected to bulge out into cells with lower pressure (Fig. 4q-q”) [53]. We constructed a weighed convexity index (wCI) to quantify convexity (Fig. 3o-q, 4o, p), and found that convexity significantly anticorrelates with number of neighbours, in agreement with qualitative observations in oryzalin-treated meristems [28], and in agreement with the pressure trends.

Altogether, our data indicate that non-random turgor pressure heterogeneity establishes in tissues with static topology (no neighbour number change) or dynamic topology (neighbour numbers change due to division). Although tissue topology is sufficient to explain pressure heterogeneity, we do not exclude a role of other biological parameters and/or sources of noise in pressure variations, as suggested by weaker correlations in experiments (Fig. 3h’, 4h’) than in simulations (Fig. 1g, h).

### Realistic mechanical models of tissue indentation are inconsistent with homogeneous pressure

Whereas the aforementioned pressure deduction is based on a model utilizing local cell shape [39], Mosca et al. [46] proposed that cell packing also contribute to the indentation stiffness. We therefore implemented realistic indentation using a thin-shell indentation finite element method (FEM) model following Mosca et al. [46]. We constructed two realistic templates (untreated- and treated-like) from confocal images; the surface was meshed and the outermost cell layer was constructed to represent the epidermis, with uniform values of cell wall thickness taken from our EM-based measurements and same cell dimensions as in confocal images; all cells were inflated by turgor pressure (Fig. 2m). We used the null hypothesis that all cells were inflated by uniform turgor pressure (2 MPa, rounded from experimental values), and performed indentations on the exact corresponding cells indented experimentally from these two templates, excluding cells at periphery of the template to avoid boundary effects. We chose indentation near the cell centre to have near-perpendicular indentation, like in experiments. We recovered values of apparent stiffness *k* that have the same magnitude as experimental values: untreated-like *k*_FEM_ = 14.9 ± 0.1 N/m (mean ± SEM, compared to *k*_AFM_ = 12.9 ± 0.2 N/m), treated-like *k*_FEM_ = 38.2 ± 1.0 N/m (compared to *k*_AFM_ = 14.5 ± 0.6 N/m). However, we found that the *k* variability differs from experiments, notably in untreated SAMs (Fig. 3k, 4k). Closer inspection revealed that cell surface mean curvature *κ*_M_, which is directly linked to *P*, also showed strong discrepancy between simulation and experiment (Fig. 3l, 4l). Accordingly, the null hypothesis of uniform turgor is not consistent with experimental data. We corrected such discrepancy based on the theory of thin shells (apparent stiffness mainly depends on these two parameters following 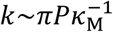, see [39]) by rescaling *κ*_M_ and *P* according to their experimental variability, and recovered the heterogeneous distribution of *k* determined by AFM (Fig. 3m, 4m). Conversely, correction of *k* and *κ*_M_ also recovered the experimentally deduced *P* distribution. This indicates that the discrepancy between FEM and AFM indentations can be fully explained by the heterogeneity in pressure (and subsequently curvature), and the contribution of cell packing to the measured variability is negligible with our depths of indentation.

### Two types of correlations between cell growth and local topology or size

Next, we monitored areal growth rate of SAM epidermal cells by time-lapse confocal microscopy. As observed previously, untreated SAMs exhibited slower growth in the centre, where stem cells reside, than the surrounding cell [54] (Fig. S5). In addition, cellular growth rate anticorrelates with neighbour number (11 SAMs, *n* = 1491 cells, *R* = −0.15, *p* = 10^−8^; Fig. 5e) and cell size (Fig. 5f; *R* = −0.33, *p* = 10^−38^), supporting previous reports that smaller cells in SAM grow faster [34,55], and suggesting that higher turgor pressure in fewer-neighboured cells associates with faster growth. In oryzalin-treated SAMs, however, the fewer-neighboured and small cells grew slower (14 SAMs, *n* = 1160 cells; Fig. 5k, neighbour number *R* = 0.20, *p* = 10^−11^; Fig. 5l, cell size *R* = 0.16, *p* = 10^−8^). This suggests that higher turgor pressure associates with either faster or slower growth depending on conditions. Although seemingly a small shift, this negative-to-positive slope change of local growth heterogeneity captures a strong qualitative inversion of growth behaviour (Fig. 5d, j). Accordingly, smaller cells expand more than larger cells in untreated SAMs, which may contribute to cell size homeostasis, a phenomenon that would be absent in oryzalin-treated SAMs.

**Figure 5.**
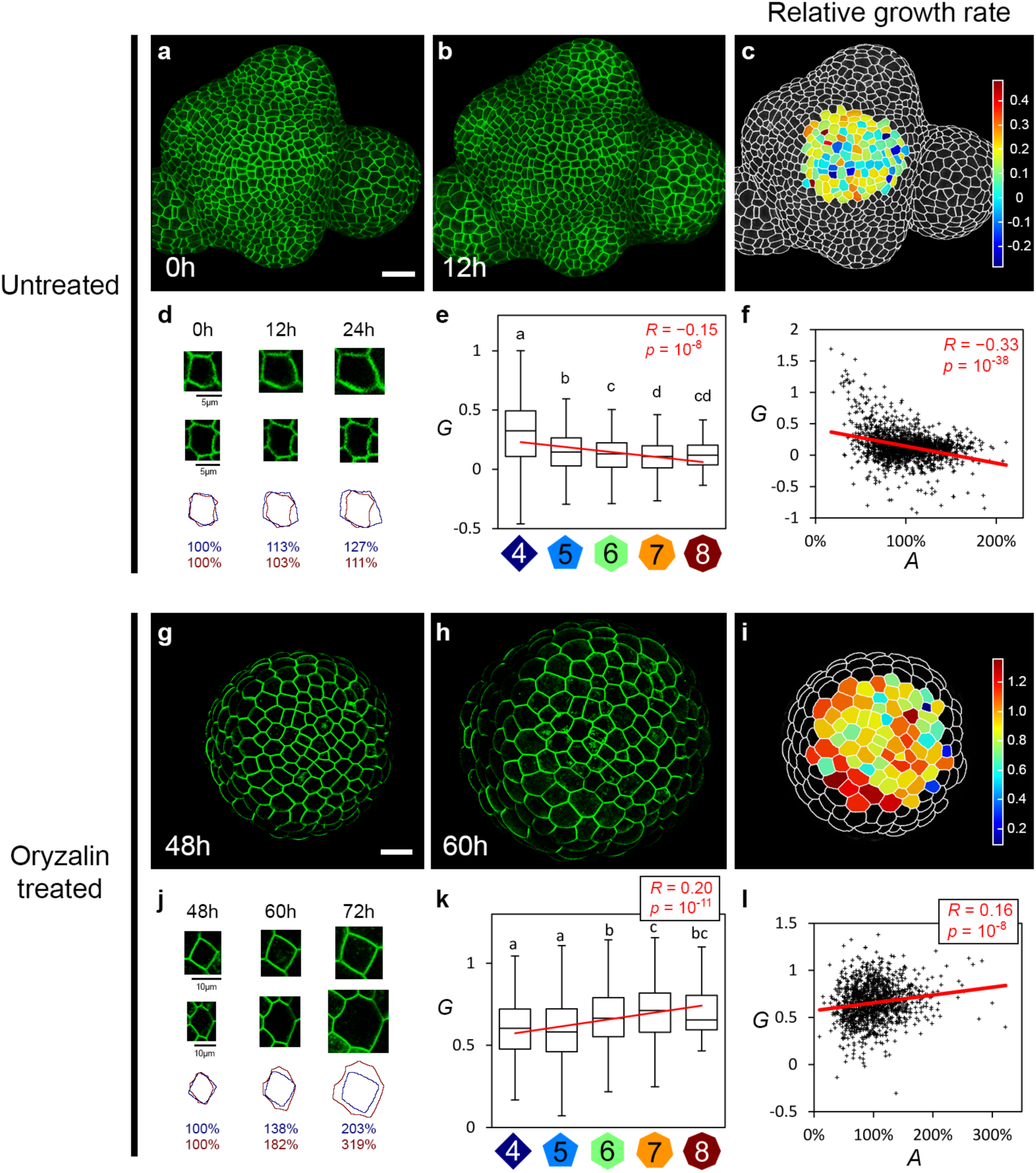
Cellular growth rate bifurcates between conditions. (**a** to **f**) Relative growth rate per day *G* of untreated SAM cells. (**g** to **l**) Cellular growth rate per day of oryzalin-treated SAM between 12-hour interval (48 and 60 hours post treatment). (a, b, g and h) Surface projections of untreated or oryzalin-treated SAM at initial time point (a and g) and 12 hours later (b and h); scale bars are 20 µm unless otherwise noted. (c and i) Heat maps of areal relative growth rate per day. (d and j) Example 4 and 8-neighbored cells during 24-hour growth, with areal normalization at initial time point. Cell contour and relative size (blue for 4-neighbored, red for 8-neighbored) depict the diverging growth trends. Scale bars are as indicated. (e, f, k and l) Box plots of relative growth rate per day *G* against cell topology *N* (e and k) and dot plots of relative growth rate per day *G* against normalized cell area *A* (f and l) (e and f, untreated 11 SAMs, *n* = 1491 cells; k and l, oryzalin-treated 14 SAMs, *n* = 1160 cells). Note that Tukey’s outliers are plotted in Figure S5 and all data are included for statistical analyses. Lowercase letters indicate statistically different populations (Student’s *t*-test, *p* < 0.05); red lines indicate linear regressions, with Pearson correlation coefficient *R* and corresponding *p*-value.

### Heterogeneity in growth rate is patterned according to the balance between wall extensibility, tissue conductivity and osmotic drive

We further explored the growth trend in our vertex model. Based on the parameter exploration on dividing mesh aimed to reproduce untreated SAM behaviour, we found both positive and negative correlations between growth rate and cell neighbour number (Fig. S2). This indicates that growth trend is sensitive to parameters that alter the balance between water flux and wall expansion (governed by the non-dimensional parameter *α*^*a*^) and to the ratio between osmotic pressure and yield pressure (*θ* = Δ*Π*/*P*^*Y*^). This can be rationalized by examining the relative growth rate *G* of a cell according to the Lockhart model (see [30]).

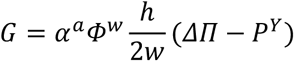

Both the prefactor *α*^*a*^ *φ*^*w*^ *h*/2*w* and the yield pressure *P*^*Y*^ decrease with cell size *R* and neighbour number N, while Δ*Π* is constant. Consequently two regimes are expected: when the osmotic drive *θ* is smaller than a threshold *θ*^*T*^, *G* is dominated by the variations of *P*^*Y*^, therefore *G* increases with cell size and with neighbour number, which corresponds to the trend in oryzalin-treated meristems; when *θ* > *θ*^*T*^, the variations of *P*^*Y*^ are negligible, so *G* follows the prefactor and decreases with cell size, which corresponds to the trend in untreated meristems. These two regimes occur whatever the value of *α*^*a*^, and the threshold value *θ*^*T*^ increases with increasing *α*^*a*^. In a tissue context, many global parameter shifts can invert growth trend through changes in these dimensionless parameters (Fig. 6k, l).

**Figure 6.**
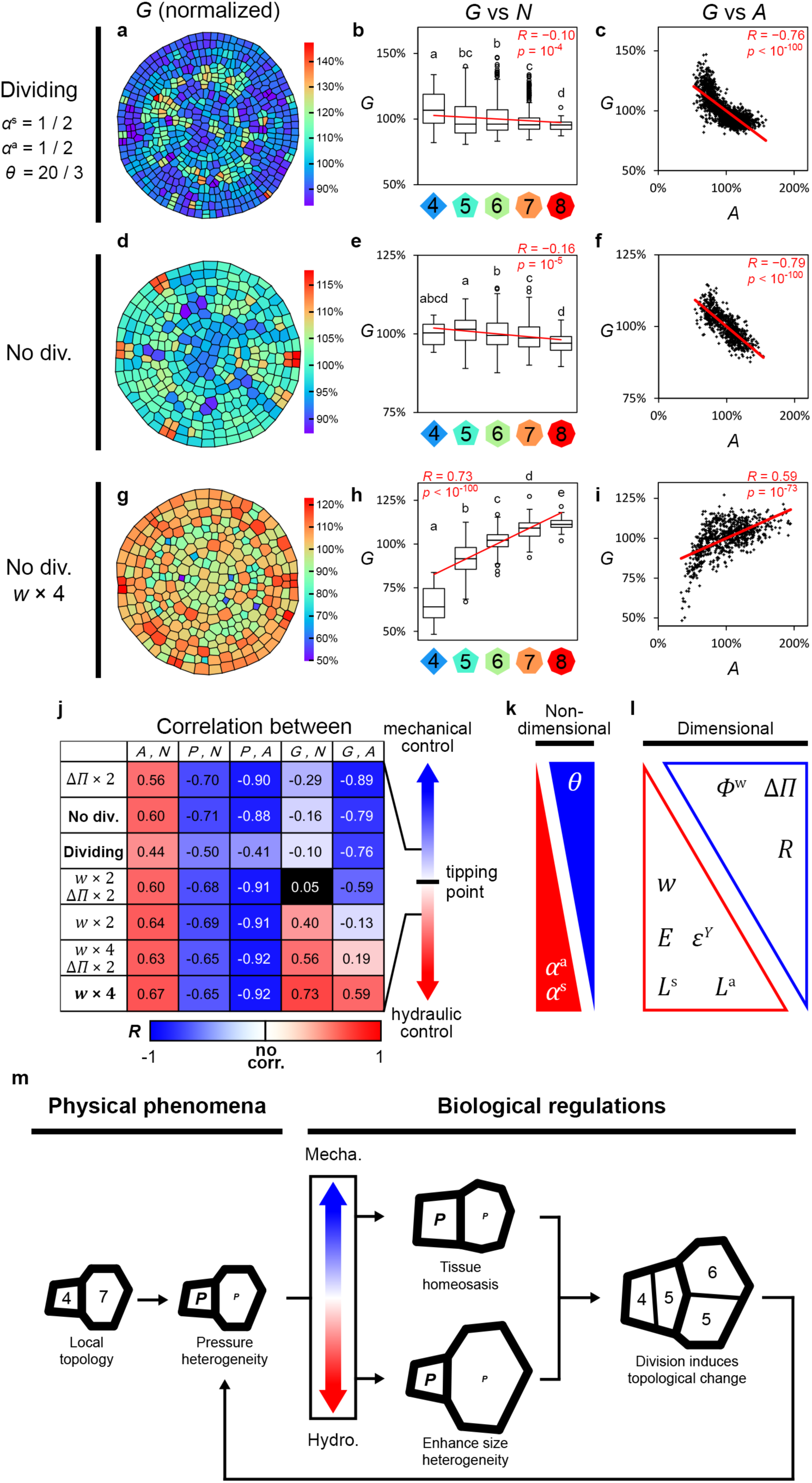
The model recapitulates the untreated and oryzalin-treated growth trends. (**a** to **i**) Relative growth rate *G* normalized by average growth rate, neighbour number *N*, and normalized cell area *A*, with reference values of dimensionless parameters (*α*^a^ = 0.5, *α*^s^ = 0.5, *θ* = 20/3): dividing simulations (a-c), non-dividing simulations (d-f), and non-dividing simulations with quadruple wall thickness *w* to mimic oryzalin treatment (g-i). (a, d, g) Heat maps of normalized areal relative growth rate. (b, e, h) Box plots of normalized relative growth rate *G* against cell topology *N*. Lowercase letters indicate statistically different populations (Student’s *t*-test, *p* < 0.05); red lines indicate linear regressions, with Pearson correlation coefficient *R* and corresponding *p*-value. (c, f, i) Dot plots of normalized relative growth rate *G* against normalized cell area *A*. For numbers of repeats see STAR Methods. (**j**) Model exploration to fit oryzalin-treated case. Colours indicate Person correlation *R*, with perfect anticorrelation as blue (*R* = −1), perfect correlation in red (*R* = 1), and insignificant correlation (*p* > 0.05) in black. *A*, normalized cell area; *N*, cell neighbour number; *P*, normalized turgor pressure; *G*, normalized relative growth rate; Δ*Π*, transmembrane osmotic pressure difference; *w*, wall thickness. (**k, l**) Influence of dimensionless (k) and dimensional parameters (l) on growth trends. Triangles indicate parameter’s influence to the mechanical– hydraulic balance. *α*^a^, dimensionless parameter for flux-wall balance; *α*^s^, dimensionless parameter for symplasmic-apoplasmic balance; *θ*, dimensionless osmotic drive. *w*, wall thickness; *E*, cell wall modulus; *ε*^*Y*^, strain threshold; *L*^a^, cross-membrane water conductivity; *L*^s^, cell-to-cell symplasmic conductivity; *Ф*^w^, wall extensibility; Δ*Π*, transmembrane osmotic pressure difference; *R*, representative cell size. (**m**) Summary of growth trend inversion according to the mechanical–hydraulic control shown in (j). The wall and flux limits correspond to untreated and oryzalin-treated SAMs, respectively.

Overall, the model retrieves untreated SAM trends if transmembrane conductivity partially limits growth (*α*^a^ not too large), symplasmic conductivity is not on par with transmembrane conductivity (*α*^s^ not too large) or if the osmotic drive is sufficient (the ratio of osmotic to yield pressure is large enough). Conservatively, we chose equal contribution by flux and wall in our model (*α*^a^ = 0.5), and increased osmotic pressure to 2 MPa, a value comparable to the experimental measurements: Figures 6a-c show that the fewer-neighboured and smaller cells grow faster (3 simulations, *n* = 1496; *G* vs *N, R* = −0.10, *p* = 10^−4^; *G* vs *A, R* = −0.76, *p* < 10^−100^).

We then attempted to reproduce oryzalin-treated behaviour, guided by the experimental observations that, besides stopping cell division, oryzalin treatment also yields higher osmotic pressure (1.6-fold) and drastically thicker cell walls (4-fold) (Fig. S3, S4). We found that both stalling division and increasing osmotic pressure failed to invert the growth trend in the model, as expected, while doubling and quadrupling wall thickness inverted the correlation of growth rate to neighbour number and cell size (Fig. 6d-j; S6), like in oryzalin-treated SAM. Combining higher osmotic pressure and thicker wall revealed that quadrupling wall thickness can robustly trigger growth trend inversion (Fig. 6j, S6).

Effectively, changing osmotic pressure and wall thickness alter the wall–flux limitation balance of the system (Fig. 6j-l). Higher osmotic pressure induces faster water influx (reduced limitation by hydraulics). The extra volume strains the walls to accumulate stress farther beyond the threshold, which is relaxed by wall yielding (growth) with extensibility as the rate limit. Meanwhile, wall thickening reduces wall stress and strain towards the threshold for expansion, effectively reducing the mechanical drive of growth and increasing the weight of water permeability in limiting growth. We do not exclude other possible parameter changes triggered by oryzalin treatment, like water conductivity and wall synthesis rate, that also contribute to the wall–flux limitation balance. Nevertheless, implementing the observed cell wall thickening in the model is sufficient to explain the observed growth rate inversion from untreated to oryzalin-treated scenario.

## DISCUSSION

In this study, we modelled the growth of a plant tissue by coupling tissue mechanics and tissue hydraulics. This generalizes previous models focusing only on mechanics [56–58]. In this model, both cell growth and turgor pressure emerge from mechanics and hydraulics. Each of these parameters can be controlled by genetic and biochemical inputs, and small uniform changes in these biological inputs can enable drastic shifts of system behaviour and the final cell size distribution (Fig. 6m). We predicted that a broad distribution of neighbour number leads to heterogeneity in cell growth and in turgor pressure, even when hydraulics has a minor contribution to the control of growth. We verified this prediction in the context of the shoot apical meristem (SAM) of *Arabidopsis thaliana*. It remains to be seen whether our results apply to other plant tissues, to animals or whether this is specific to the SAM.

In order to test our predictions and deduce pressure in the SAM, we combined a recently developed indentation-based approach [40] with FEM-based realistic mechanical models of indentation [46]. We found values of turgor in the range 1-3 MPa range, higher than the range 0.2-1 MPa generally measured in plant tissues [18], though values of up to 5 MPa were measured in guard cells [48]. For instance, the Arabidopsis root epidermis has a turgor of about 0.4 MPa, as measured with the pressure probe [59]; the Arabidopsis leaf epidermis has a turgor of about 1-2 MPa, as deduced form indentation and mechanical modelling [60]. Accordingly, we speculate that the SAM function might require relatively high turgor. Another specificity of the SAM could be a relatively low transmembrane conductivity, as most aquaporin (channel protein allowing rapid transmembrane water flux [61]) isoforms had significantly lower expression in inflorescence than in other fast growing tissues like stem and root [62] (with a reduction up to ∼ 80%). Nevertheless, a wide range of water conductivity can produce pressure heterogeneity, including very high conductivity (i.e. *α*^a^ = 0.9 [36]), and the agreement between model predictions and experimental measurements suggest that tissue hydraulics has a (possibly small) contribution to limiting growth in the SAM. Similarly, altered expression of the aquaporin PIP2;1 delays the emergence of lateral roots [63], also hinting to a developmental role for hydraulic conductivity. Altogether, we propose that tissue mechanics and hydraulics act in parallel with the established genetic regulations in SAM cell growth.

We found that both cell growth and turgor pressure are heterogeneous in the SAM. It has already been reported that smaller cells [55] or the smaller of two sister cells [34] grow faster in the SAM of untreated and NPA-treated plants, respectively. We found the same trend in untreated SAM, with an inversion in oryzalin-treated SAM. Such inversion is not irrelevant to normal development: for example, a growth variability switch is observed during sepal development, where clones of smaller and often fewer-neighboured cells grow faster in young sepals then slower in older sepals, effectively shifting from homogenizing to amplifying cell size variability [64]. In untreated SAM, small cells grow faster than big cells, possibly contributing to tissue homeostasis. Finally, irrespective of the conditions, we find that turgor pressure is smaller in big cells in experiments, which might contribute to reducing mechanical stress in the cell wall in these cells [65]; this would act in parallel with the mechanism proposed in the context of leaf epidermal cells, based on cells adopting puzzle shapes that limit cell wall stress [65].

Our results point towards a link from cell topology (number of neighbours) to cell mechanical status. This might also be relevant to animal epithelia [29], though this appears unexplored experimentally. Feedbacks from cell mechanics to cell topology are more established: cell division and thus number of neighbours can be oriented by mechanical stress in animals and in plants [66–69]. Since tissue topology is highly conserved in many biological systems [70], we propose that pressure heterogeneity may emerge in compact tissues with polygonal cells [63] and non-instantaneous water movement, due to the adjustment to reconcile local mechanical and hydraulic conditions.

Finally, we note that heterogeneous patterns may not always be stochastic [71]. The emergent heterogeneity of local growth and hydrostatic pressure is coupled with the characteristic yet dynamic tissue topology [51,52] (Fig. 6m), all based on stringent rules and likely underlies morphogenesis in compact tissues. With the discovery of many cell-size-dependent transcripts [3,72], our model proposes another source for non-random variability in a tissue.

## STAR Methods

### Plant materials, treatments and growth conditions

*Arabidopsis thaliana GFP-LTi6b* (ecotype WS-4) reporter line and ecotype Col-0 was used[73]. Untreated inflorescence meristems were obtained from soil-grown plants, first in short-day (8 h light 20°C / 16 h dark 19°C cycle) for 3 to 4 weeks then transferred to long-day (16 h light 20°C / 8 h dark 19°C cycle) for 1 to 2 weeks to synchronize bolting. Oryzalin-treated inflorescence meristems were obtained from plants grown on custom-made Arabidopsis medium [74] (Duchefa) supplemented with 1% agar-agar (Merck) and 10 µM N-1-naphthylphthalamic acid (NPA, Sigma-Aldrich/Merck) for 3 weeks. Pin-formed inflorescence meristems from NPA medium were immersed in 10 µg/mL oryzalin (Sigma-Aldrich/Merck) twice (3 h duration, 24 h interval) [28]. For mechanical measurements and time-lapse confocal imaging, meristems were mounted on Arabidopsis apex culture medium (ACM) [74] with 2% agarose and 0.1% plant preservation mixture (PPM, Plant Cell Technology) to prevent contamination, and cultivated in long-day condition.

### Atomic force microscopy

Untreated meristems (dissected, with most late stage-2 floral primordia removed to prevent blocking of the cantilever) and oryzalin-treated meristems were mounted on ACM (2% agarose, 0.1% PPM) the night before. Drops of 2% low melting agarose (Duchefa) were applied around the lower parts of meristems for mechanical stabilization. For oryzalin-treated meristems, 72 h post-treatment meristems were measured.

AFM indentations were performed as in Beauzamy et al., 2015 [40]. Specifically, a BioScope Catalyst model AFM (Bruker) operated with the NanoScope software (version 9.1, Bruker), under a MacroFluo optical epifluorescence macroscope (Leica), was used. All measurements were done with customized 0.8 µm diameter spherical probes mounted on silicon cantilevers of 42 N/m spring constant (SD-Sphere-NCH-S-10, Nanosensors). Cantilever deflection sensitivity was calibrated against a clean sapphire wafer submerged in water before each session.

Meristems were submerged in water during AFM measurements. PeakForce QNM mode was used to record sample surface topography and cell contours (aided by the stiffness difference between periclinal and anticlinal cell walls on DMT modulus maps) in overlapping square tiles of 30×30 to 50×50 µm^2^ (128×128 pixels). Force curves were obtained by the point-and-shoot mode of the NanoScope software, with at least 3 locations chosen near the barycentre of each cell, and 3 consecutive indentations per location, making at least 9 force curves per cell. Approximately 10 µN maximum force was applied during each indentation, corresponding to approximately 1 µm indentation depth.

For hyperosmotic treatments, oryzalin-treated meristems were mounted in Petri-dishes on Patafix (UHU), then the gap between Patafix and sample base was quickly sealed with bio-compatible glue Reprorubber-Thin Pour (Flexbar) for stabilization. After the glue solidified (less than 2 minutes), samples were submerged in liquid ACM containing 0.1% PPM. Samples were first measured in liquid ACM (plus 0.1% PPM), then submersion medium was changed to ACM plus desired concentration of NaCl (plus 0.1% PPM) by first rinsing with 3∼ 5 mL target solution, then soaked in target solution for 5 minutes before AFM measurements. Each new measurement per solution change took around 30 minutes.

### Force curve analysis

Turgor pressure was determined as previously reported [40]. Specifically, cell wall elastic modulus (Hertzian model, 1∼ 10% maximal indentation force) and cell apparent stiffness (linear, 75∼ 99% maximal indentation force) were retrieved from each force curve by the NanoScope Analysis software (version 1.5, Bruker). Quality of force curves were checked empirically and by the fit coefficient of determination *r*^*2*^ > 0.99. Cells with only low quality force curves were not analysed. Cell surface curvatures (mean and Gaussian) were estimated from AFM topographic images, with the curvature radii fitted to the long and short axes of each cell. Turgor pressure was further deduced from each force curve (100 iterations) with the electron-microscopy-determined cell wall thickness 190 nm for untreated and 740 nm for oryzalin-treated SAMs, and cell-specific turgor pressure is retrieved by averaging all turgor deductions per cell.

For cell registration, confocal stacks of each meristem were obtained prior to AFM measurements by an LSM 700 confocal (Carl Zeiss). Surface projection of *GFP-LTi6b* signal was generated by the software MerryProj [75], then rescaled and rotated (affine transformation) to overlay the AFM image tiles. The resulting surface projection image was used to generate cell contour image of the whole meristemic surface using morphological segmentation plugin [76] for the software ImageJ (https://fiji.sc/), while the relative positions of each AFM indentation location is then registered onto the cell contour image, along with cellular geometrical and topological analyses, using the NanoIndentation plugin (version alpha) for ImageJ [77].

Since each meristem had different turgor pressure range, cellular turgor pressure was normalized to the average of each meristem for comparing cell-to-cell turgor pressure heterogeneity without meristem-specific effects.

### Electron microscopy

For serial block-face imaging SEM (SBF-SEM), plants were grown in vitro on medium containing the auxin transport inhibitor NPA (Naphtalene Phtalamic Acid) to generate stems with naked meristems and were locally treated for 48h with the microtubule depolymerizing drug oryzalin (Sigma) in lanolin at a concentration of 2 µg/µl [78]. These plantlets were subsequently taken off the inhibitor and left to regenerate for 48h on normal Arabidopsis medium. Meristems with young organ primordia were fixed in 0.5% glutaraldehyde (in demineralized water), from 25% Sigma stock in Microscopy Facility lab. The plantlets were left at room temp for 2h in an eppendorf and rinsed 1x in water before post fixation and dehydrating and embedding in Spurr’s epoxy as described in [79]. The samples were then sectioned and viewed in a Zeiss Merlin SEM [79].

For standard transmission electron microscopy fixed meristems of soil grown plants were embedded in Spurr’s resin and sectioned before viewing in a Jeol 2100F (at the Centre Technologique des Microstructures, UCBL, Lyon).

### Time-lapse confocal microscopy

Untreated (dissected) and oryzalin-treated meristems were mounted and grown on ACM with 0.8% agarose and 0.1% PPM for live imaging. Confocal stacks were taken on an LSM 700 confocal microscope (Carl Zeiss) operated with the ZEN 2010 software (version 6.0, Carl Zeiss), using a W N-Achroplan 40x/0.75 M27 water immersion objective, and on a TCS SP8 confocal microscope (Leica) operated with the Leica Application Suite X software (version 3.5, Leica), using a Fluotar VISIR 25x/0.95 water immersion lens. GFP was excited at 488 nm and emission detected between 415 – 735 nm. Stacks have resolution of 1028×1028 pixels, with resolution ranging between 3.2 to 4.4 pixels/µm; Z steps were between 0.5 and 0.85µm.

For hyperosmotic treatments, meristems were mounted in Petri-dishes on Patafix (UHU), then submerged in liquid ACM containing 0.1% PPM. Samples were first imaged in liquid ACM (plus 0.1% PPM), then submersion medium was changed to ACM plus desired concentration of NaCl (plus 0.1% PPM) by first rinsing with 3∼ 5 mL target solution, then soaked in target solution for 5 minutes before imaging. Because of the reduced signal in hyperosmotic solutions, possibly due to the altered refractive index, stronger gain was used to reach comparable signal intensity.

### Image processing and geometric analysis

3D shell mesh and surface projection of untreated meristems were generated from confocal stacks using the level set method (LSM) addon [80] for the software MorphoGraphX (MGX version 1.0) [81]. For oryzalin-treated meristems, 2D surface projections were generated by MerryProj [75] and imported into MGX for further processing. Projected images were segmented using watershed method after manual seeding, and cell lineage between time points was manually assigned in the meristem proper. To limit Z distortion and biases due to change in inclination of the surface, which may affect analysis accuracy, only cells within 20° of inclination angle from the highest position of the SAM were included. A custom-made Python script was used to trace cell lineage between multiple time points and determine cell topology based on the anticlinal wall number exported from MGX. Areal relative exponential growth rate per hour was calculated as:

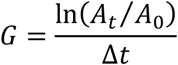

where Δ*t* is time interval in hours, *A*_*0*_ is original cellular area at time *t*_*0*_, and *A*_*t*_ is final area at time *t*_*0*_ *+* Δ*t*. Cells undergone topological changes (i.e. divided cells and cells adjacent to new division planes) during the acquisition were not included in the growth analyses. To analyse variation, cell-specific growth rate was further co-aligning by the median per SAM then stretching the distribution to the average first and third quartile positions to each data point.

To enhance the confocal images in hyperosmotic solution, anisotropic diffusion filter [82] was re-implemented and applied to the raw images with the following parameters (specifications see http://cbp-domu-forge.ens-lyon.fr/redmine/projects/anifilters/wiki): *K* = 0.3, *σ* = 5, *γ* = 0.9, *D* = 10, 50 iterations.

For figure panels, brightness and contract of confocal images were linearly enhanced for better visual. To synchronize panel shape and size, black background with no relevant information was cropped from or added to the edge of the panels.

### Finite element method (FEM) indentation simulation

Inflation and indentation simulations were performed with the same procedure adopted from Mosca et al., 2017 [46]. For each meristem, a surface projection of the L1 anticlinal walls obtained from confocal microscopy was segmented and extruded to the average anticlinal wall height as observed in the template (5 µm for untreated and 10µm for oryzalin-treated) with the CellMaker plugin for MorphoGraphX [81]. This generates a multicellular template made of triangular membrane elements. The extruded meshes keep the average overall organ curvature.

In order to give the extruded cells a more realistic rest curvature (in the unpressurized state), both meshes were pre-inflated with a small pressure (0.05 MPa for untreated and 0.15 MPa oryzalin-treated SAM) and saved as the rest configuration. Both meshes were refined around the indentation points to increase the accuracy of the indentation simulations. The untreated mesh used in the indentation simulation and curvature analysis has an average edge size of 0.1 µm near the indentation points and increases to 0.5 µm far away from those points. The oryzalin-treated mesh has instead an average edge size of 0.2 µm and 1 µm near and far from the indentation the indentation areas, respectively.

The cell wall was modelled as a Saint-Venant Kirchhoff material represented by membrane elements with zero transversal stress and mathematically prescribed thickness [46]. The two templates were assigned the following material properties:

- isotropic material, 200 MPa Young’s modulus, 0.3 Poisson ratio;
- cell wall thickness of 0.19 µm and 0.74 µm for the untreated and oryzalin-treated meristem, following experimental data.

The templates were inflated from the rest configuration with a turgor pressure of 2 MPa inside each cell for both meshes, until the forces equilibrium was reached (sum of the forces squared). With the chosen parameters, the total cell height after inflation is similar to the experimental observation from the confocal stacks for the referring templates (6.8 µm for untreated and 14.5 µm for oryzalin-treated). Afterwards the bottom of the template (bottom anticlinal cell walls) was blocked in all space degrees of freedom to simulate the presence of the supporting inner tissue during the indentation process.

The indentation is modelled as in [46] and is performed in the global z-direction as given by the confocal images. The indentation process is performed on each cell independently, where the specific indentation point is chosen to be close to the uppermost anticlinal wall location. This reproduces the AFM indentation modality. As in analysis of AFM experiments, the stiffness of the untreated cell was computed as the slope of the indentation curve at 0.5 µm indentation depth, with the reaction forces on the indenter between 0.3 and 0.7 µm depth used for slope computation. For the oryzalin-treated, given the thicker walls, the indentations were deeper and the stiffness was computed around 1 µm indentation depth (reaction forces between 0.8 and 1.2 µm).

For both meshes, a refinement analysis was performed to verify the accuracy of the simulation results. The results are reported in Supplementary Table 1. We considered the variation of the coarse oryzalin mesh so small to justify using it for the AFM and curvature comparison analysis, while we preferred using the refined mesh for the untreated to reduce the error due to mesh resolution.

**Supplementary Table 1.** FEM refinement analysis. Coarse mesh edge size near indentation; refined mesh average edge size near indentation; relative MD, relative mean absolute difference; sd, standard deviation.

| Mesh | coarse size | refined size | Relative MD | sd of relative MD |
| --- | --- | --- | --- | --- |
| Untreated | 0.2 $\mu\text{m}$ | 0.1 $\mu\text{m}$ | 3% | 2.4% |
| Oryzalin-treated | 0.4 $\mu\text{m}$ | 0.2 $\mu\text{m}$ | 2% | 1.3% |

### Statistical analysis

Data were processed using Excel 2000 (Microsoft). All Tukey box plots depict the first, second (median) and third quartiles of data distribution, with whiskers marking the lowest/highest data within 1.5 interquartile ranges (IQR) of the lower/upper quartiles. Tukey’s outliers are depicted as small circles outside the whiskers. Values like turgor pressure, cell area and growth rate were normalized to the average per meristem. After normalization, every cell was considered as one biological sample, and all linear regressions and Pearson correlations were performed on whole datasets. For simulations, cells on the edge of the mesh were not analysed due to border effect. Extremely rare polygon classes (i.e. triangle and nonagon) were not shown on the box plots in the main figures but were included in linear regression and Pearson correlation tests and were plotted in Figure S5. Kolmogorov–Smirnov test was used to distinguish cell neighbour number distribution (significance level *α* = 0.05).

### Mechanical-hydraulic modelling

#### Summary

We build a vertex-based model of plant tissues at cellular level that couples osmosis-driven hydraulic fluxes between cells and from apoplast with a fixed water potential, and cell wall mechanics which resists and grows under tension. Turgor and growth rate heterogeneities emerge from this coupling and from the heterogeneities in cells sizes and topology (number of neighbours).

We consider a collection of *N* polygonal cells *i* = 1, …, *N* that form a mesh; this mesh evolves with the appearance of new cells because of cell division. The walls between cells are discretized into one or several segments. Given the topology, the mesh is fully characterized by the position of the vertices. The walls are given a height *h* and a thickness *w*.

#### Cell wall rheology

The cell walls are modelled as a visco-elasto-plastic material, which would be equivalent to the Ortega model [20] in the case of an elongating cell. Let *σ*_*k*_ be the stress of a wall segment *k*; the constitutive law writes 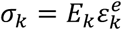 where *E*_*k*_ is the elastic modulus and 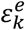 is the elastic deformation of the wall. Let *l*_*k*_ be the length of segment *k*, the rate of change of 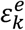 is given by:

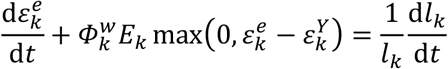

where 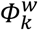 is the extensibility and 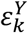 is the yield deformation of segment *k*. Equivalently, we could define a yield stress.

#### Mechanical equilibrium

Let *P*_*i*_ be the turgor pressure in each cell *i*. The tissue being at every moment in a quasi-static equilibrium, pressure forces on wall edges and elastic forces within walls balance exactly at each vertex *v*:

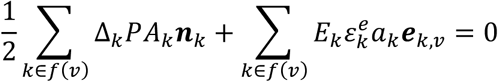

Where *f*(*v*) is the set of walls adjacent to junction *v*, and 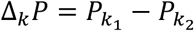 is the pressure jump across wall face *k*, with *k*_*1*_ < *k*_*2*_ as indices of the cells separated by face *k, A*_*k*_ = *hl*_*k*_ is the area of the face *k* on which pressure is exerted, ***n***_*k*_ is the normal vector to face *k*, oriented from cell *k*_*1*_ to cell *k*_*2*_, and *a*_*k*_ = *hw* is the cross-section of the face, on which the elastic stress is exerted; finally, ***e***_*k,v*_ is the unit vector in the direction of face *k*, oriented from junction *v* to the other end of face *k*. In the case of a single cylindrical cell for which growth is restricted to its principal direction, the model is equivalent to the Lockhart-Ortega model.

#### Fluxes

For each cell *i*, the apoplasmic pathway is represented as a flux 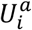 (in volume per time unit) from the apoplast of constant water potential Ψ^*a*^ through a perfectly semi-permeable membrane: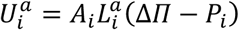, where *A*_*i*_ is the area of each cell in contact with the apoplast, 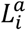 is the corresponding water conductivity, Δ*Π* = *π*_*i*_ + Ψ^*a*^ is assumed constant, and *π*_*i*_ is the osmotic pressure of cell *i*.

The symplasmic pathway corresponds to flows that occur through plasmodesmata, channels between cells that convey both water and solutes. The symplasmic flows thus only depend on turgor pressure difference. Let *L*_*ij*_ be the symplasmic water conductivity corresponding to the interface between two neighbour cells *i* and *j*, and *A*_*ij*_ their contact area, both assumed symmetric: *L*_*ij*_ = *L*_*ji*_ and *A*_*ij*_ = *A*_*ji*_. The symplasmic flux 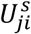 (in volume per time unit) from cell *j* to *i* is defined by:

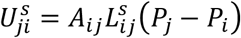

Finally, the total water flux for cell *i* is the sum of the apoplasmic flux 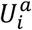 and the symplasmic fluxes 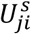 with all its neighbors, so that its volume variation can be expressed as:

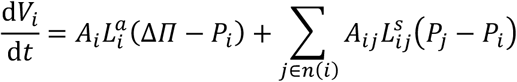

where *n*(*i*) is the set of neighbours of cell *i*.

#### Cell division

We implemented the Willis-Refahi rule [34], in which the division volume is given by *v*_0_ = *f V*_*b*_ + *μ*_*b*_(2 - *f* + *Z*), where *f* = 0.5, *V*_*b*_ is the volume at birth, *μ* _*b*_ = 3.31 is the mean birth volume and *Z* is a Gaussian noise with zero mean and 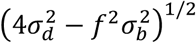 standard deviation, with *σ*_*b*_ = 0.2 and *σ*_*d*_ = 0.1.

#### Numerical resolution

In the Lockhart-Ortega model, the compatibility between wall elongation and cell volume increase is automatically enforced through the geometrical constraint of unidirectional growth that leads to equal relative growth rate of the cell and strain rate of the walls. In our multicellular model, this equality is no longer true. Instead, the lengths *l*(*X*) of the edges and the volumes *V*(*X*) of the cells are expressed as functions of the positions *X* of the vertices; then, given an initial position *X* of the vertices and elastic deformation *ε*^*e*^ of the edges, the equations of wall rheology, mechanical equilibrium, and water fluxes form a closed set of equations with respect to the unknowns *X, P*, and *ε*^*e*^ that allow to predict their evolution.

To give an idea of the mathematical complexity of the problem, one may consider the following example: in a connected tissue, if one cell is stretched and forced to increase its volume, an equal volume of water has to enter the cell, either from the apoplastic compartment or the neighbour cells. In the latter case, pressure should drop in the neighbour cells, which should attract water from their own neighbours, and this could propagate to further cells depending on the geometry of the tissue and the effective parameters. Volume and therefore positions of the vertices could be also affected. Finally, one can see that the interaction between hydraulics and mechanics implies long range interactions where pressure plays a key role.

We developed an original algorithm and implemented it in an in-house code, where at each time step, the mechanical equilibrium is resolved under constraints on the cell volume (from the water fluxes), and constraints on the cells edges (from the rheological law of the walls). This was implemented in Python, using the open source libraries NumPy, SciPy, and the Topomesh class from the OpenAlea project (http://openalea.gforge.inria.fr/doc/vplants/container/doc/html/container/openalea_container_topomesh_ref.html). This algorithm is described in more detail in a separate publication [30].

The computations were run on a computer with a 3.6GHz Intel Xeon E5 processor, 64 GB of RAM, and running Linux Debian Stretch. The typical computing time was one week for each computation.

#### Parameterization of the model

The reference values of the parameters were chosen based on the literature or on our experiments (main text Table 1), except for *α*^s^, for which no data is available and was conservatively ascribed an intermediate value of 0.5. Model behaviour was explored by changing non-dimensional parameter values as explained in the main text.

#### Procedure for the computations

We first run in parallel three computations with cell division to around 300 cells. To mimic untreated case, simulations continue until around 600 cells. To mimic the oryzalin treatment, the current states of the “untreated” computations at around 300 cells are used as initial conditions for the oryzalin case: division is stopped, and we run computations either with the same effective parameters, or with some parameters modified so that the behaviour of the oryzalin treated meristems is recovered (see supplementary table 2); the computations are run until the total volume has been multiplied by three from this initial state. Typical runtime for one parameter set is one week.

**Supplementary table 2.**
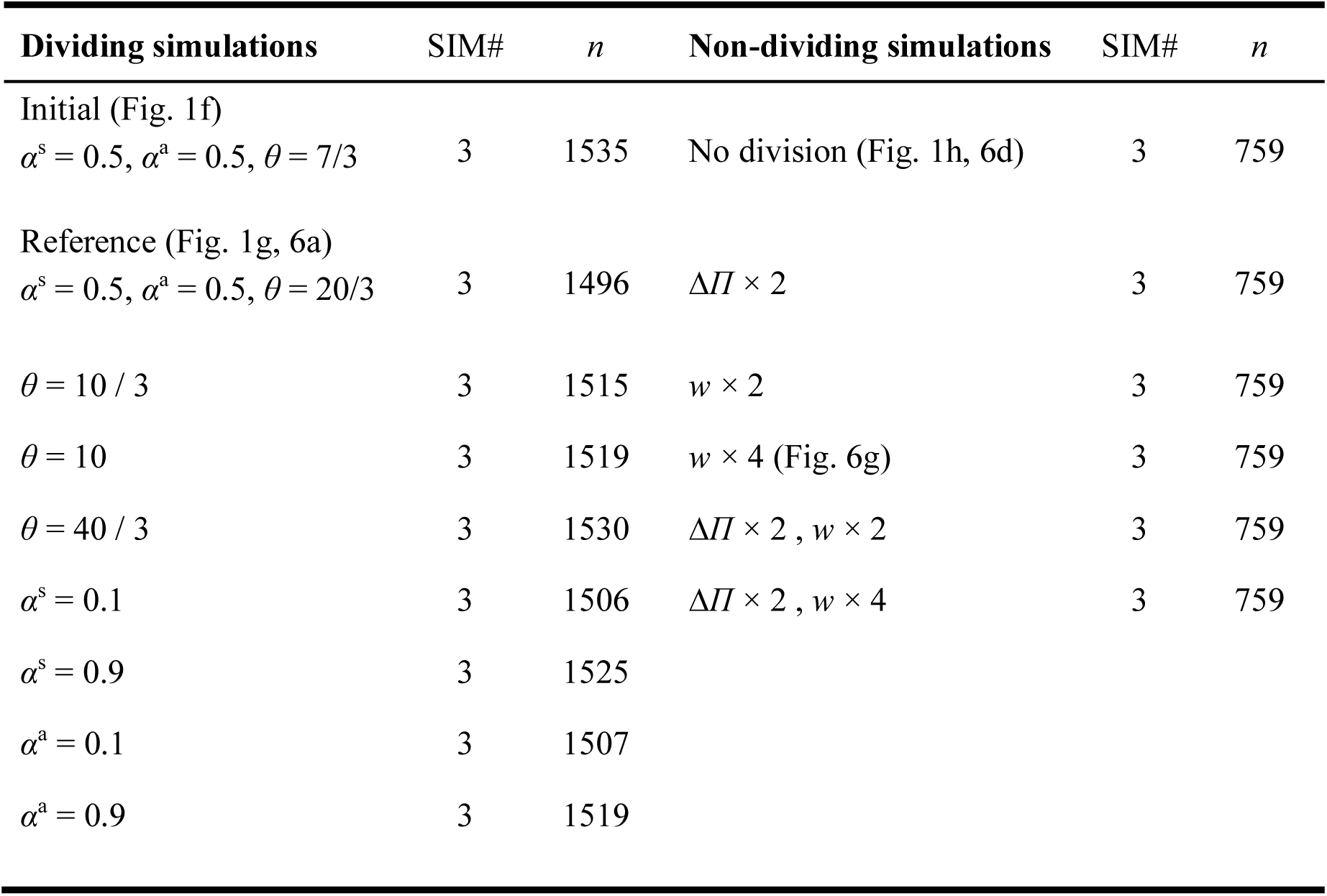
Parameter modification and simulation repeats. Parameters are systemically changed, with the reference parameter set *α*^s^ = 0.5, *α*^a^ = 0.5, *θ* = 20/3 for exploration in untreated simulations and for mimicking oryzalin in non-dividing simulations. Annotations are as described above. SIM#, number of simulations; *n*, total cell number at last time point without boundary cells.

## Acknowledgements

We thank P. Bolland and A. Lacroix for plant care, G. Cloarec for help with TEM, L. Beauzamy for AFM training, C. Mollier for help in estimating confocal technical errors, A. Kiss for help with anisotropic filtering parameterization, G. Cerutti for help in implementing the division algorithm and in segmentation, R. Smith for help in segmentation and mesh generation for the indentation simulations, and V. Battu, F. Zhao and C. Galvan-Ampudia for providing plant materials. We acknowledge the contribution of the PLATIM facility of SFR Biosciences (UMS3444/CNRS, US8/Inserm, ENS de Lyon, UCBL) for AFM and confocal microscopy, and of Centre Technologique des Microstructures (UCBL, Lyon) for electron microscopy. This work was supported by a fellowship from Institut Universitaire de France and an ERC Starting Grant “PhyMorph” to A.B. (ERC-2012-StG-307387), an EMBO Long-term Fellowship to Y.L. (EMBO ALTF 168-2015), and an Agropolis Foundation grant (MecaFruit3D) to I.C. and C.G.

The authors declare no competing financial interests.

## Author contributions

This study was initiated by A.B. Y.L. and A.B. designed the experiments. Y.L. executed AFM and confocal microscopy, acquired experimental data except EM, and analysed experimental and simulation data. J.T. performed electron microscopy. V.M and M.D. wrote scripts to facilitate experimental data analysis. I.C. and C.G. designed physical model of tissue growth.I.C. implemented the model, ran simulations, optimized model parameters and analysed simulation data. G.M. designed, ran, and analysed indentation simulations. Y.L. and A.B. contributed to the design of simulations. Y.L., I.C., V.M., G.M., C.G. and A.B. contributed to data interpretation. Y.L. and A.B. wrote the manuscript with inputs from the other authors.

## Data availability

All materials, scripts and datasets generated and analysed during the current study are available from the corresponding authors upon reasonable request.

## Supplementary figures

**Figure S1.**
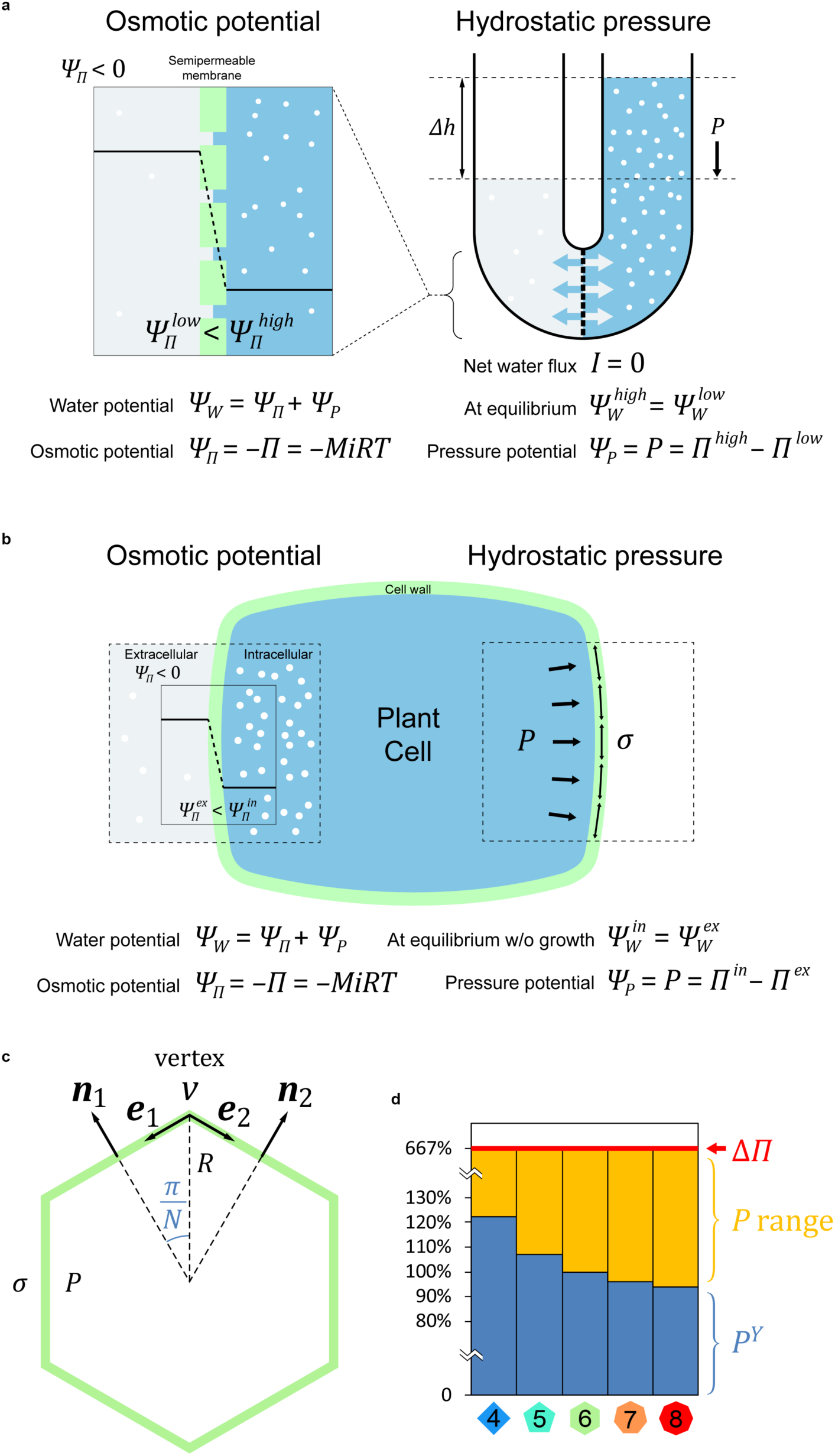
Osmotic pressure and hydrostatic pressure. (**a**) Osmosis occurs when solutions of different osmotic pressure *Π*, due to different solute concentration (depicted by the density of white dots), are separated by a semipermeable membrane. Difference of osmotic potential *Ψ*_*Π*_ (depicted by horizontal black lines, always negative in solutions) across the membrane dictates that solvent flows from high to low potential compartment (low to high solute concentration). In a U-shaped tube setup (right), osmosis may stop before the two compartments reach the same concentration, as the extra volume (in fact height, Δ*h*) in the higher-concentration compartment exerts a hydrostatic pressure *P* due to gravity that pushes the solvent back. *P* increases until reaching the same value as Δ*Π* at equilibrium where the total water potential *Ψ*_*W*_ is equal on both sides, and the net solvent flux *I* = 0 (indicated by the opposite blue arrows). *M*, solute molar concentration; *i*, van t’Hoff index of solute that disassociates; *R*, ideal gas constant; *T*, absolute temperature. (**b**) Osmosis in a plant cell, where gravity is neglected for its small size. Because of the rigid cell wall that restricts the cell volume, hydrostatic pressure *P*, alias turgor pressure, builds up alongside cell wall tension *σ* to counterbalance the difference of osmotic potential *Ψ*_*Π*_, until *P* = Δ*Π*. (**c**) Geometrical extension of Lockhart model for a single polygonal cell. *σ*, cell wall stress; *P*, turgor pressure; *R*, cell radius; *N*, polygon class; v, vertex; ***n***_**1**_ and ***n***_**2**_; normal vector; ***e***_**1**_ and ***e***_**2**_, tangent vector. (**d**) Half of the threshold pressure of a polygonal cells base on 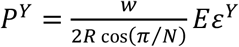. For growing cells, turgor pressure *P* falls between the transmembrane osmotic pressure Δ*Π* and pressure threshold *P*^*Y*^.

**Figure S2.**
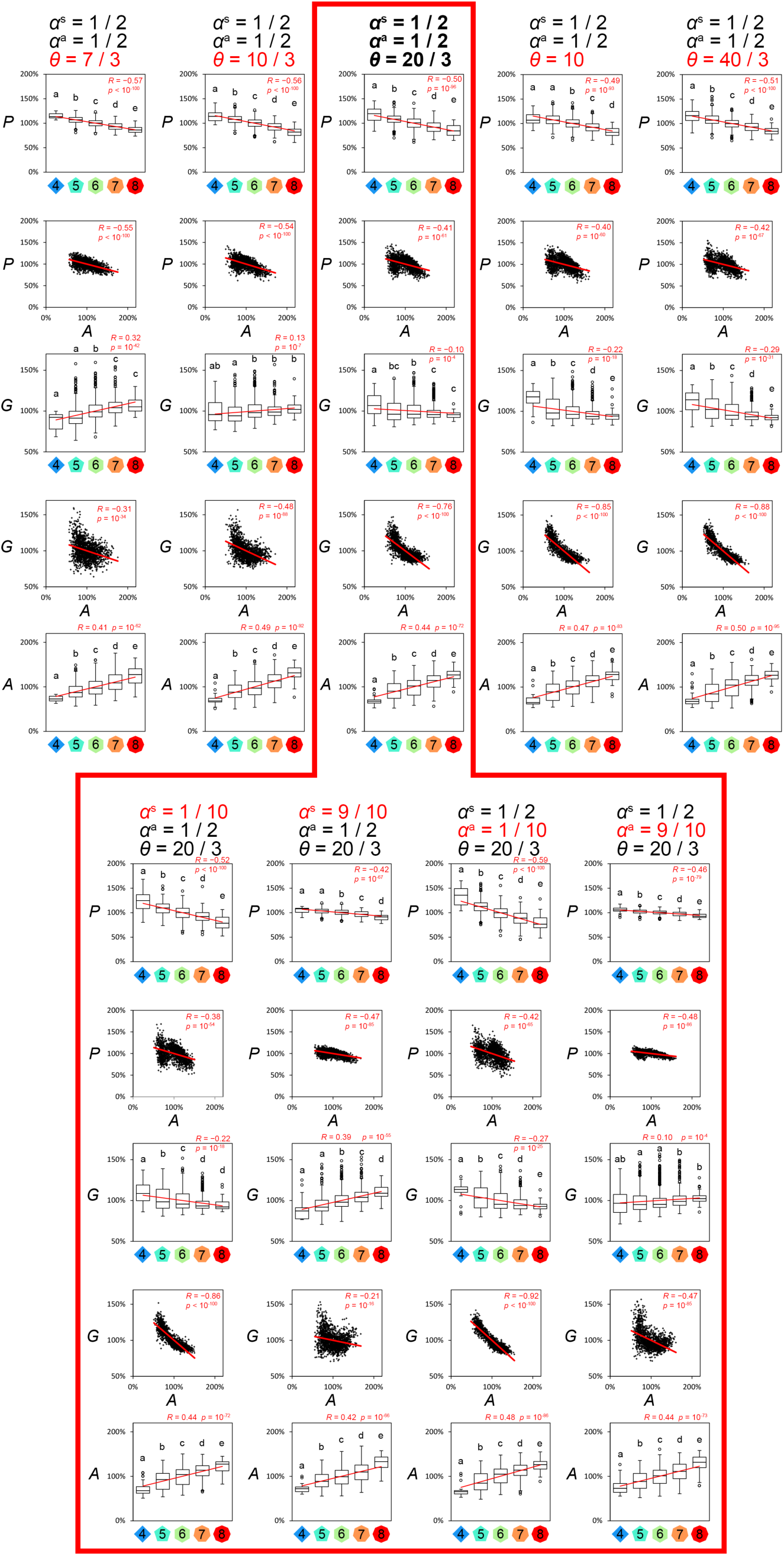
Simulations for vertex model parameter exploration. Model parameters were systemically changed. *α*_a_, dimensionless parameter determining the weight of the limitation of growth by transmembrane permeability to the combined limitation of growth by cell wall extensibility and transmembrane permeability; *α*_s_, dimensionless parameter determining the weight of intercellular water conductivity over total (intercellular and transmembrane) conductivity; *s*, cell size scale at division relative to the initial setting. *P*, normalized turgor pressure, *A*, normalized cell area; *G*, relative growth rate per hour. Polygons indicate cell neighbour number. Red lines indicate linear regressions, with Pearson correlation coefficient *R* and corresponding *p*-value.

**Figure S3.**
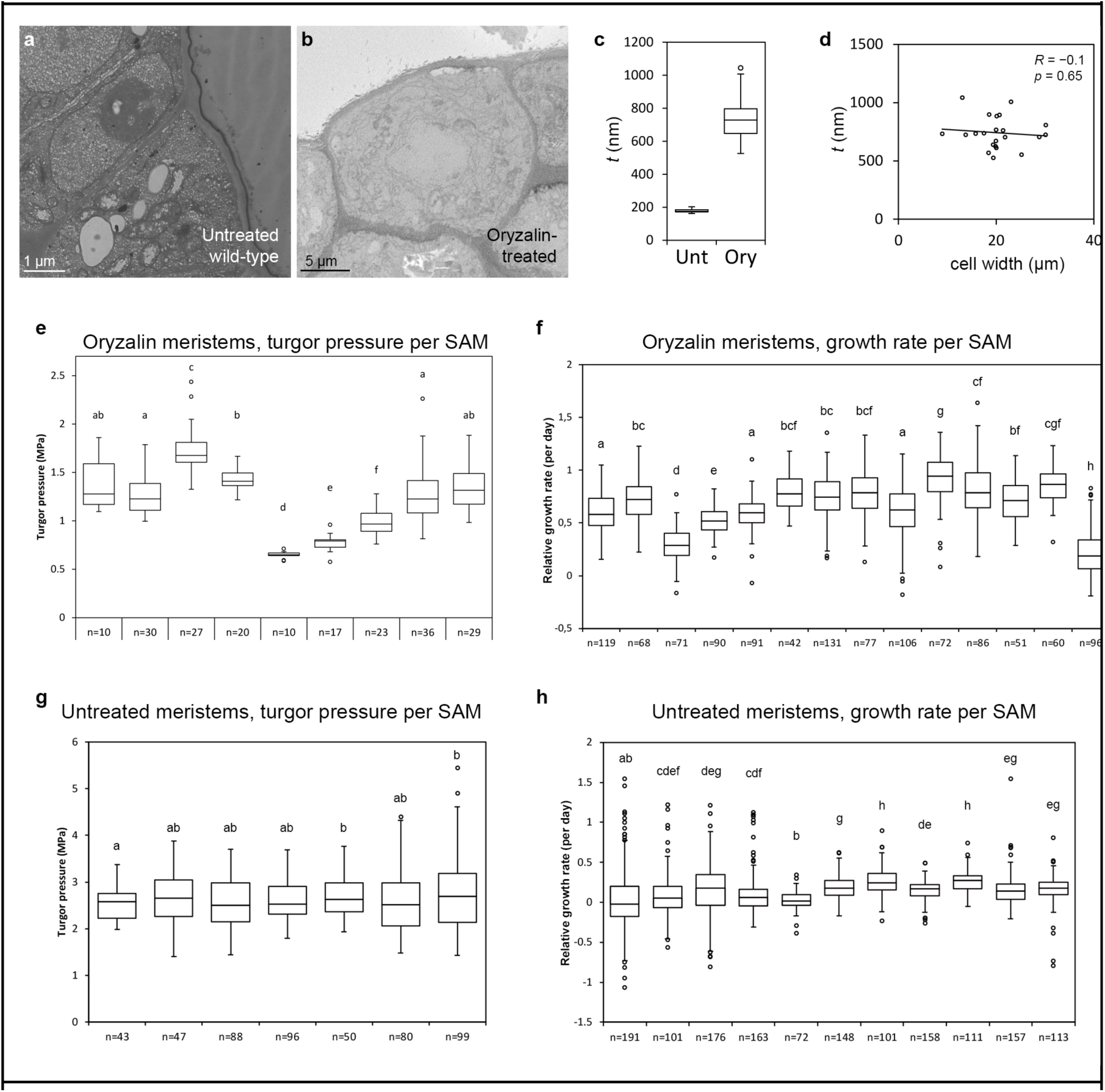
Cell wall thickness and individual SAM measurements of turgor pressure and growth rate. (**a** and **b**) TEM and serial block-face imaging SEM images of untreated (a) and oryzalin-treated meristem (b). (**c**) Boxplots of surface wall thickness of untreated (Unt) and oryzalin-treated (Ory) meristems. (**d**) Surface wall thickness does not correlate to cell width in oryzalin-treated meristems. (**e-h**) Oryzalin-treated SAMs showed highly variable turgor pressure and growth rate per SAM, while untreated samples show little variation; *n* is cell number measured per meristem. Circles are Tukey’s outliers; lowercase letters indicate statistically different populations (Student’s *t*-test, *p* < 0.05). (g) The first five untreated meristems measured by AFM were of ecotype WS-4, while the last two are of Col-0.

**Figure S4.**
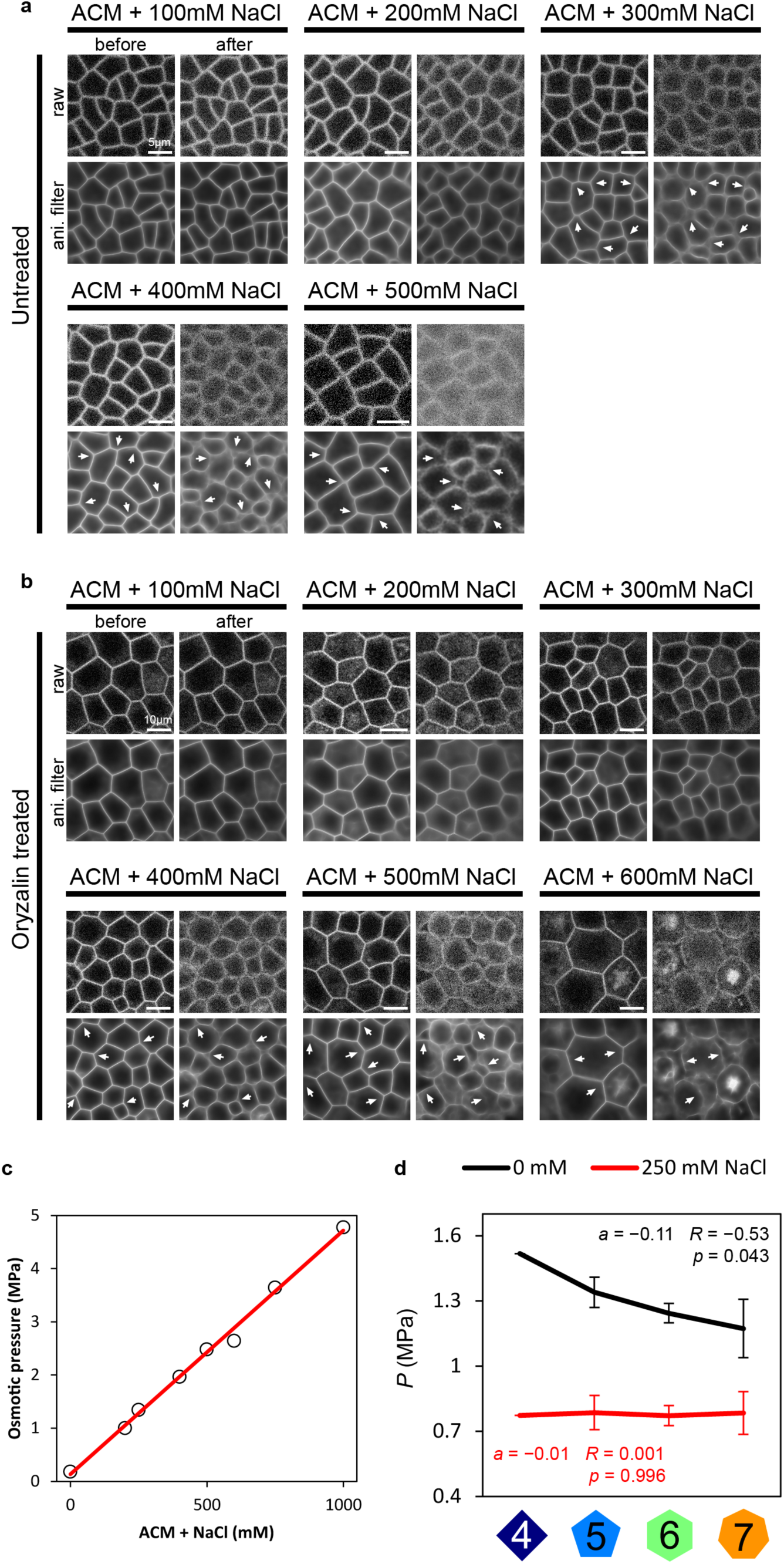
Hyperosmotic treatments on SAMs. (**a** and **b**) Confocal images of untreated (a) and oryzalin-treated (b) meristems submerged in liquid apex culture medium (ACM) before and after the supplementation of NaCl. The increased osmolality resulted in blurred images (each upper-right subpanel), possibly due to altered refraction limit, which required anisotropic filtering to enhance the trajectory of cell contours. Out of 8 untreated meristems, half showed incipient plasmolsysis at 200 mM NaCl (∼0.4 Osm) and the other half at 300 mM (∼0.6 Osm). For 7 oryzalin-treated meristems, incipient plasmolysis occurred among a broader osmolarity (2 meristems at 300 mM, 3 meristems at 400 mM, 2 meristems at 500 mM). Arrows points to noticeable membrane detachment and rounding, indicating plasmolysis. (**c**) Osmolarity calibration by osmometer measurements (*r*^2^ > 0.99). (**d**) Moderate hyperosmotic treatment decreases AFM-determined turgor pressure, and reduces turgor heterogeneity plotted against cell neighbour number (linear regression slope *a*, Pearson correlation coefficient *R* and corresponding *p*-value).

**Figure S5.**
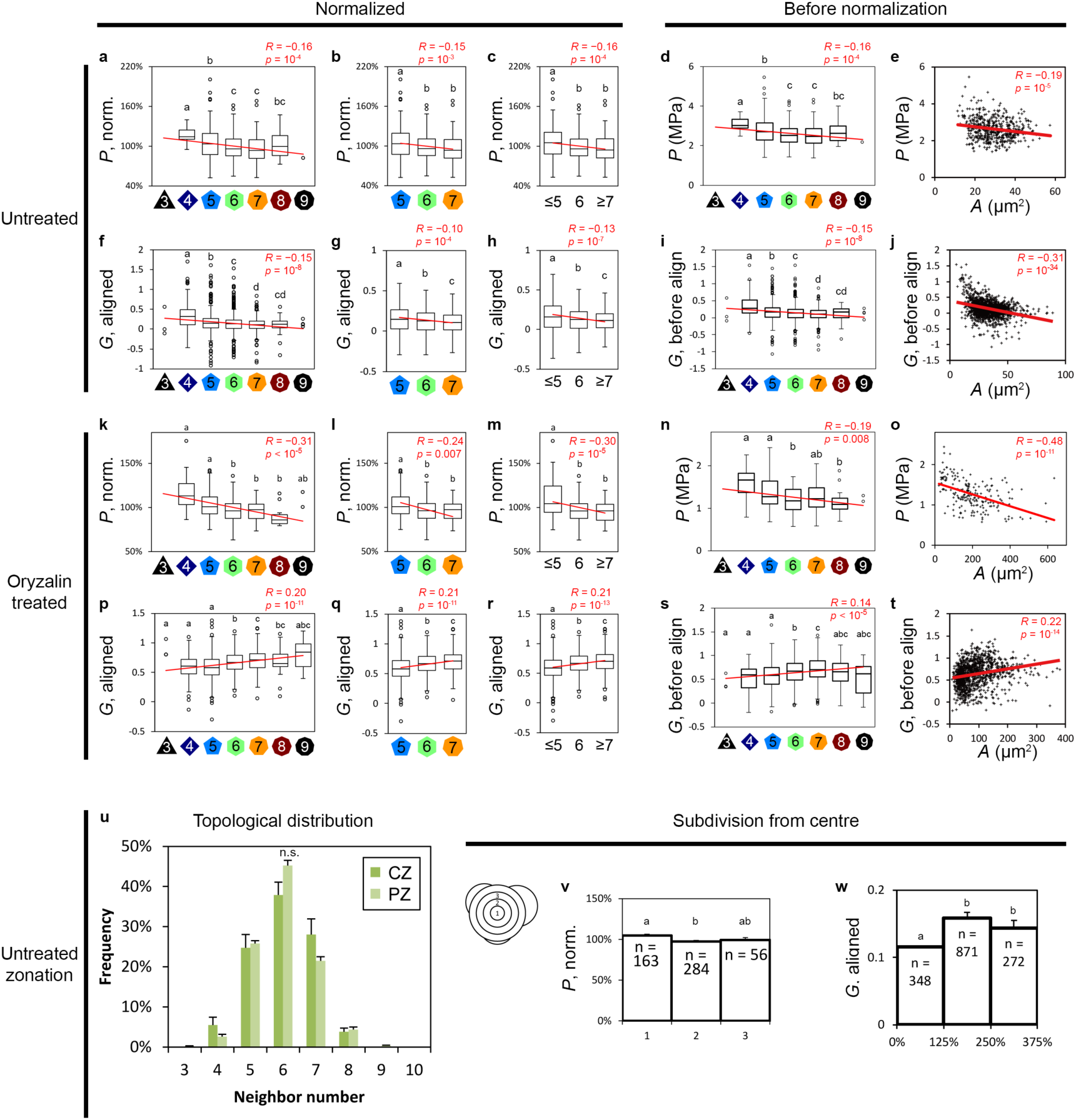
Pressure and growth rate before and after normalization, and regional measurements. (**a** to **j**) Normalized and raw turgor pressure *P* and growth rate *G* of untreated meristems plotted against cell neighbour number *N* and cell area *A*. (**k** to **t**) Normalized and raw turgor pressure *P* and growth rate *G* of oryzalin-treated meristems plotted against cell neighbour number and cell area. (a, f, k, p) Normalized *P* and co-aligned *G* against *N* plots complete with outliers and the rare 3- and 9-neighboured cells, same data as in Figure 3 and 4. (b, g, l, q) Same normalized data with only the major polygonal classes (*N* = 5, 6, 7), showing that the correlations do not depend on the rare polygonal classes at the ends of distributions. (c, h, m, r) Same normalized data were binned for cells with 5 or fewer neighbours (≤5), cells with 6 neighbours and cells with 7 or more neighbours (≥7), showing the same trends as unbinned data. (d, i, n, s) Raw data of *P* and *G* before normalization / co-alignment against neighbour number; (e, j, o, t) raw data of *P* and *G* before normalization / co-alignment against unnormalized cell area, all showing the same trend as normalized data. Red lines are linear regression line, with Pearson correlation coefficient *R* and corresponding *p*-value. Lower case letters denote statistically significant groups (Student’s *t*-test *p* < 0.05). (**u**) Frequency of *N*-neighboured cells in central zone (CZ) and peripheral zone (PZ) shows no significant difference (Kolmogorov–Smirnov test, confidence level *α* = 0.05, *D*_*n,m*_ < *D*_*α*_). (**v, w**) Normalized turgor pressure and co-aligned growth rate in subregions of SAM; *n* is cell number, error bars indicate standard error of mean, lower case letters denote statistically significant groups (Student’s *t*-test *p* < 0.05).

**Figure S6.**
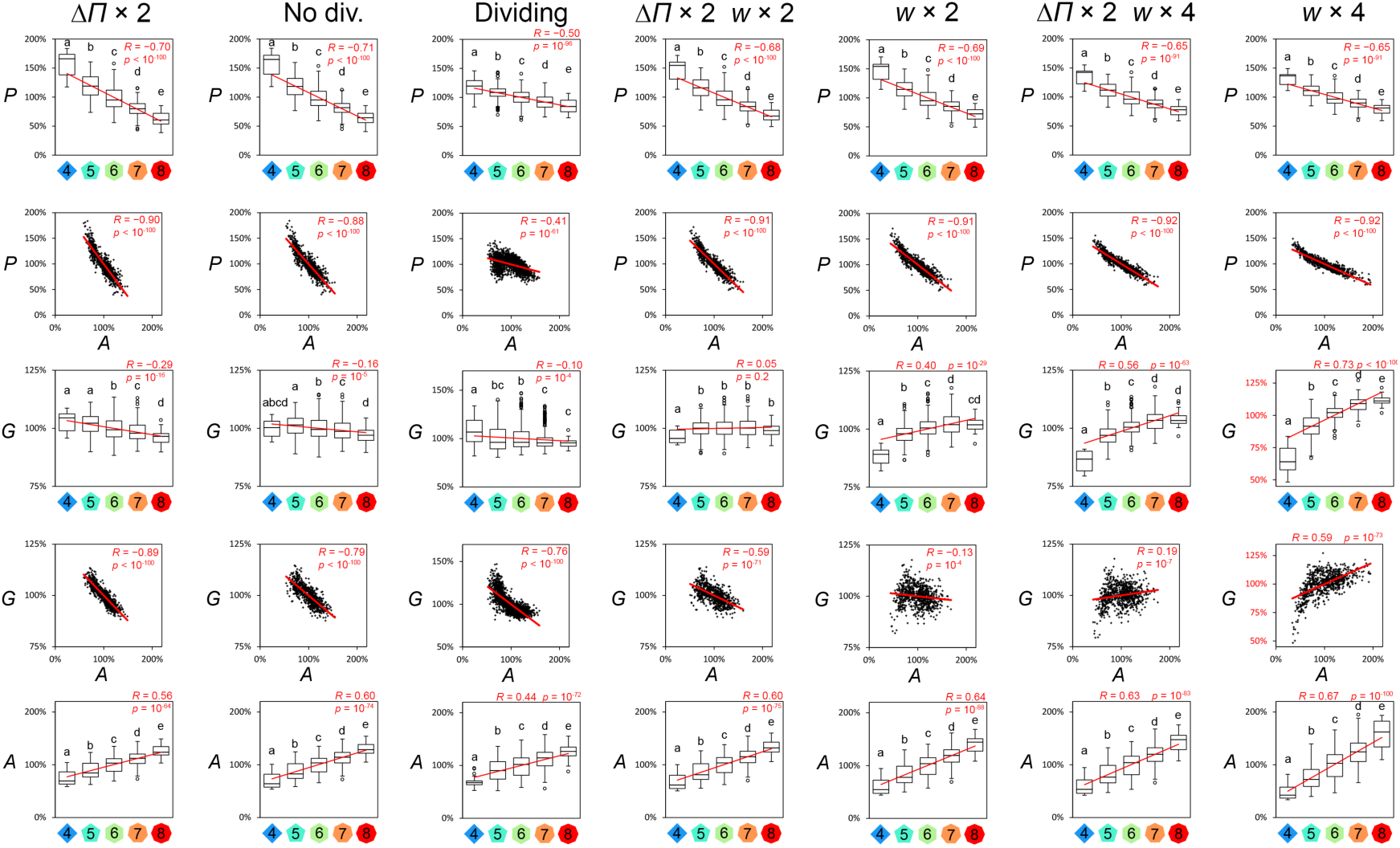
Simulations to recapitulate oryzalin-treated SAM. Based on the reference parameter set (*α*^s^ = 0.5, *α*^a^ = 0.5, *θ* = 20/3), which represents the untreated-like behaviour with cell divisions (Dividing), oryzalin-treated-like behaviour was further fitted by stopping cell division (No div.) and subsequently changing osmotic pressure Δ*Π* and cell wall thickness *w*, based on experimental observations. *P*, normalized turgor pressure, *A*, normalized cell area; *G*, normalized relative growth rate. Polygons indicate cell neighbour number. Red lines indicate linear regressions, with Pearson correlation coefficient *R* and corresponding *p*-value.

## Notes

#### Summary of Updates

We performed new simulations with experimentally-based parameters. We reordered the manuscript so that to make the conclusions clearer.

## References

1. Hong, L.; Dumond, M.; Zhu, M.; Tsugawa, S.; Li, C.-B.; Boudaoud, A.; Hamant, O.; Roeder, A.H.K. Heterogeneity and Robustness in Plant Morphogenesis: From Cells to Organs. Annu. Rev. Plant Biol. 2018, 69, 469–495.

2. Kamimoto, K.; Kaneko, K.; Kok, C.Y.-Y.; Okada, H.; Miyajima, A.; Itoh, T. Heterogeneity and stochastic growth regulation of biliary epithelial cells dictate dynamic epithelial tissue remodeling. eLife 2016, 5.

3. Ietswaart, R.; Rosa, S.; Wu, Z.; Dean, C.; Howard, M. Cell-Size-Dependent Transcription of FLC and Its Antisense Long Non-coding RNA COOLAIR Explain Cell-to-Cell Expression Variation. Cell Syst. 2017, 4, 622-635.e9.

4. Long, Y.; Stahl, Y.; Weidtkamp-Peters, S.; Postma, M.; Zhou, W.; Goedhart, J.; Sánchez-Pérez, M.-I.; Gadella, T.W.J.; Simon, R.; Scheres, B.; et al. In vivo FRET-FLIM reveals cell-type-specific protein interactions in Arabidopsis roots. Nature 2017, 548, 97–102.

5. Chubb, J.R. Symmetry breaking in development and stochastic gene expression. Wiley Interdiscip. Rev. Dev. Biol. 2017, 6, e284.

6. Donati, G.; Watt, F.M. Stem Cell Heterogeneity and Plasticity in Epithelia. Cell Stem Cell 2015, 16, 465–476.

7. Gerdes, M.J.; Sood, A.; Sevinsky, C.; Pris, A.D.; Zavodszky, M.I.; Ginty, F. Emerging Understanding of Multiscale Tumor Heterogeneity. Front. Oncol. 2014, 4.

8. Eldar, A.; Elowitz, M.B. Functional roles for noise in genetic circuits. Nature 2010, 467, 167–173.

9. Watanabe, K.; Umeda, T.; Niwa, K.; Naguro, I.; Ichijo, H. A PP6-ASK3 Module Coordinates the Bidirectional Cell Volume Regulation under Osmotic Stress. Cell Rep. 2018, 22, 2809–2817.

10. Xie, K.; Yang, Y.; Jiang, H. Controlling Cellular Volume via Mechanical and Physical Properties of Substrate. Biophys. J. 2018, 114, 675–687.

11. Guo, M.; Pegoraro, A.F.; Mao, A.; Zhou, E.H.; Arany, P.R.; Han, Y.; Burnette, D.T.; Jensen, M.H.; Kasza, K.E.; Moore, J.R.; et al. Cell volume change through water efflux impacts cell stiffness and stem cell fate. Proc. Natl. Acad. Sci. 2017, 114, E8618–E8627.

12. Dumais, J.; Forterre, Y. “Vegetable Dynamicks”: The Role of Water in Plant Movements. Annu. Rev. Fluid Mech. 2012, 44, 453–478.

13. Stewart, M.P.; Helenius, J.; Toyoda, Y.; Ramanathan, S.P.; Muller, D.J.; Hyman, A.A. Hydrostatic pressure and the actomyosin cortex drive mitotic cell rounding. Nature 2011, 469, 226–230.

14. Montel, F.; Delarue, M.; Elgeti, J.; Malaquin, L.; Basan, M.; Risler, T.; Cabane, B.; Vignjevic, D.; Prost, J.; Cappello, G.; et al. Stress clamp experiments on multicellular tumor spheroids. Phys. Rev. Lett. 2011, 107, 188102.

15. Rojas, E.R.; Huang, K.C. Regulation of microbial growth by turgor pressure. Curr. Opin. Microbiol. 2017, 42, 62–70.

16. Zerzour, R.; Kroeger, J.; Geitmann, A. Polar growth in pollen tubes is associated with spatially confined dynamic changes in cell mechanical properties. Dev. Biol. 2009, 334, 437–446.

17. Lopez, R.; Badel, E.; Peraudeau, S.; Leblanc-Fournier, N.; Beaujard, F.; Julien, J.-L.; Cochard, H.; Moulia, B. Tree shoot bending generates hydraulic pressure pulses: a new long-distance signal? J. Exp. Bot. 2014, 65, 1997–2008.

18. Beauzamy, L.; Nakayama, N.; Boudaoud, A. Flowers under pressure: ins and outs of turgor regulation in development. Ann. Bot. 2014, mcu187.

19. Feng, W.; Lindner, H.; Robbins, N.E.; Dinneny, J.R. Growing Out of Stress: The Role of Cell- and Organ-Scale Growth Control in Plant Water-Stress Responses. Plant Cell 2016, 28, 1769–1782.

20. Ortega, J.K. Augmented growth equation for cell wall expansion. Plant Physiol. 1985, 79, 318–320.

21. Kroeger, J.H.; Zerzour, R.; Geitmann, A. Regulator or driving force? The role of turgor pressure in oscillatory plant cell growth. PloS One 2011, 6, e18549.

22. Sager, R.E.; Lee, J.-Y. Plasmodesmata at a glance. J. Cell Sci. 2018, 131, jcs209346.

23. Willmer, C.M.; Sexton, R. Stomata and plasmodesmata. Protoplasma 1979, 100, 113–124.

24. Wille, A.C.; Lucas, W.J. Ultrastructural and histochemical studies on guard cells. Planta 1984, 160, 129–142.

25. Ruan, Y.L.; Llewellyn, D.J.; Furbank, R.T. The control of single-celled cotton fiber elongation by developmentally reversible gating of plasmodesmata and coordinated expression of sucrose and K+ transporters and expansin. Plant Cell 2001, 13, 47–60.

26. Rygol, J.; Pritchard, J.; Zhu, J.J.; Tomos, A.D.; Zimmermann, U. Transpiration Induces Radial Turgor Pressure Gradients in Wheat and Maize Roots. Plant Physiol. 1993, 103, 493–500.

27. Robbins, N.E.; Dinneny, J.R. Growth is required for perception of water availability to pattern root branches in plants. Proc. Natl. Acad. Sci. 2018, 115, E822–E831.

28. Corson, F.; Hamant, O.; Bohn, S.; Traas, J.; Boudaoud, A.; Couder, Y. Turning a plant tissue into a living cell froth through isotropic growth. Proc. Natl. Acad. Sci. U. S. A. 2009, 106, 8453–8458.

29. Ishihara, S.; Sugimura, K.; Cox, S.J.; Bonnet, I.; Bellaïche, Y.; Graner, F. Comparative study of non-invasive force and stress inference methods in tissue. Eur. Phys. J. E Soft Matter 2013, 36, 9859.

30. Cheddadi, I.; Génard, M.; Bertin, N.; Godin, C. Coupling water fluxes with cell wall mechanics in a multicellular model of plant development. PLOS Comput. Biol. 2019, 15, e1007121.

31. Lucas, W.J.; Ham, B.-K.; Kim, J.-Y. Plasmodesmata - bridging the gap between neighboring plant cells. Trends Cell Biol. 2009, 19, 495–503.

32. Kumar, N.M.; Gilula, N.B. The Gap Junction Communication Channel. Cell 1996, 84, 381–388.

33. McLean, P.F.; Cooley, L. Protein Equilibration through Somatic Ring Canals in Drosophila. Science 2013, 340.

34. Willis, L.; Refahi, Y.; Wightman, R.; Landrein, B.; Teles, J.; Huang, K.C.; Meyerowitz, E.M.; Jönsson, H. Cell size and growth regulation in the Arabidopsis thaliana apical stem cell niche. Proc. Natl. Acad. Sci. U. S. A. 2016, 113, E8238–E8246.

35. Boyer, J.S.; Cavalieri, A.J.; Schulze, E.-D. Control of the rate of cell enlargement: Excision, wall relaxation, and growth-induced water potentials. Planta 1985, 163, 527–543.

36. Cosgrove, D.J. Cell Wall Yield Properties of Growing Tissue: Evaluation by in Vivo Stress Relaxation. Plant Physiol. 1985, 78, 347–356.

37. Ortega, J.K.E. Dimensionless Numbers to Analyze Expansive Growth Processes. Plants 2019, 8, 17.

38. Vella, D.; Ajdari, A.; Vaziri, A.; Boudaoud, A. The indentation of pressurized elastic shells: from polymeric capsules to yeast cells. J. R. Soc. Interface 2012, 9, 448–455.

39. Vella, D.; Ajdari, A.; Vaziri, A.; Boudaoud, A. Indentation of Ellipsoidal and Cylindrical Elastic Shells. Phys. Rev. Lett. 2012, 109, 144302.

40. Beauzamy, L.; Derr, J.; Boudaoud, A. Quantifying Hydrostatic Pressure in Plant Cells by Using Indentation with an Atomic Force Microscope. Biophys. J. 2015, 108, 2448–2456.

41. Beauzamy, L.; Louveaux, M.; Hamant, O.; Boudaoud, A. Mechanically, the Shoot Apical Meristem of Arabidopsis Behaves like a Shell Inflated by a Pressure of About 1 MPa. Front. Plant Sci. 2015, 6.

42. Malgat, R.; Faure, F.; Boudaoud, A. A Mechanical Model to Interpret Cell-Scale Indentation Experiments on Plant Tissues in Terms of Cell Wall Elasticity and Turgor Pressure. Front. Plant Sci. 2016, 7.

43. Lazarus, A.; Florijn, H.C.B.; Reis, P.M. Geometry-induced rigidity in nonspherical pressurized elastic shells. Phys. Rev. Lett. 2012, 109, 144301.

44. Kierzkowski, D.; Nakayama, N.; Routier-Kierzkowska, A.-L.; Weber, A.; Bayer, E.; Schorderet, M.; Reinhardt, D.; Kuhlemeier, C.; Smith, R.S. Elastic domains regulate growth and organogenesis in the plant shoot apical meristem. Science 2012, 335, 1096–1099.

45. Milani, P.; Gholamirad, M.; Traas, J.; Arnéodo, A.; Boudaoud, A.; Argoul, F.; Hamant, O. In vivo analysis of local wall stiffness at the shoot apical meristem in Arabidopsis using atomic force microscopy. Plant J. Cell Mol. Biol. 2011, 67, 1116–1123.

46. Mosca, G.; Sapala, A.; Strauss, S.; Routier-Kierzkowska, A.-L.; Smith, R.S. On the micro-indentation of plant cells in a tissue context. Phys. Biol. 2017, 14, 015003.

47. Hong, L.; Dumond, M.; Tsugawa, S.; Sapala, A.; Routier-Kierzkowska, A.-L.; Zhou, Y.; Chen, C.; Kiss, A.; Zhu, M.; Hamant, O.; et al. Variable Cell Growth Yields Reproducible Organ Development through Spatiotemporal Averaging. Dev. Cell 2016, 38, 15–32.

48. Yi, H.; Chen, Y.; Wang, J.Z.; Puri, V.M.; Anderson, C.T. The stomatal flexoskeleton: how the biomechanics of guard cell walls animate an elastic pressure vessel. J. Exp. Bot. 2019, 70, 3561–3572.

49. Knoblauch, J.; Mullendore, D.L.; Jensen, K.H.; Knoblauch, M. Pico gauges for minimally invasive intracellular hydrostatic pressure measurements. Plant Physiol. 2014, pp.114.245746.

50. Nakayama, N.; Smith, R.S.; Mandel, T.; Robinson, S.; Kimura, S.; Boudaoud, A.; Kuhlemeier, C. Mechanical Regulation of Auxin-Mediated Growth. Curr. Biol. 2012, 22, 1468–1476.

51. Lewis, F.T. The correlation between cell division and the shapes and sizes of prismatic cells in the epidermis of cucumis. Anat. Rec. 1928, 38, 341–376.

52. Gibson, W.T.; Veldhuis, J.H.; Rubinstein, B.; Cartwright, H.N.; Perrimon, N.; Brodland, G.W.; Nagpal, R.; Gibson, M.C. Control of the mitotic cleavage plane by local epithelial topology. Cell 2011, 144, 427–438.

53. Hamant, O.; Heisler, M.G.; Jönsson, H.; Krupinski, P.; Uyttewaal, M.; Bokov, P.; Corson, F.; Sahlin, P.; Boudaoud, A.; Meyerowitz, E.M.; et al. Developmental Patterning by Mechanical Signals in Arabidopsis. Science 2008, 322, 1650–1655.

54. Kwiatkowska, D. Surface growth at the reproductive shoot apex of Arabidopsis thaliana pin-formed 1 and wild type. J. Exp. Bot. 2004, 55, 1021–1032.

55. Serrano-Mislata, A.; Schiessl, K.; Sablowski, R. Active Control of Cell Size Generates Spatial Detail during Plant Organogenesis. Curr. Biol. 2015, 25, 2991–2996.

56. Ali, O.; Mirabet, V.; Godin, C.; Traas, J. Physical models of plant development. Annu. Rev. Cell Dev. Biol. 2014, 30, 59–78.

57. Dumond, M.; Boudaoud, A. Physical Models of Plant Morphogenesis. In Mathematical Modelling in Plant Biology; Morris, R.J., Ed.; Springer International Publishing: Cham, 2018; pp. 1–14 ISBN 978-3-319-99070-5.

58. Alt, S.; Ganguly, P.; Salbreux, G. Vertex models: from cell mechanics to tissue morphogenesis. Philos. Trans. R. Soc. B Biol. Sci. 2017, 372.

59. Dyson, R.J.; Vizcay-Barrena, G.; Band, L.R.; Fernandes, A.N.; French, A.P.; Fozard, J.A.; Hodgman, T.C.; Kenobi, K.; Pridmore, T.P.; Stout, M.; et al. Mechanical modelling quantifies the functional importance of outer tissue layers during root elongation and bending. New Phytol. 2014, 202, 1212–1222.

60. Forouzesh, E.; Goel, A.; Mackenzie, S.A.; Turner, J.A. In vivo extraction of Arabidopsis cell turgor pressure using nanoindentation in conjunction with finite element modeling. Plant J. 2013, 73, 509–520.

61. Maurel, C.; Boursiac, Y.; Luu, D.-T.; Santoni, V.; Shahzad, Z.; Verdoucq, L. Aquaporins in Plants. Physiol. Rev. 2015, 95, 1321–1358.

62. Klepikova, A.V.; Kasianov, A.S.; Gerasimov, E.S.; Logacheva, M.D.; Penin, A.A. A high resolution map of the Arabidopsis thaliana developmental transcriptome based on RNA-seq profiling. Plant J. Cell Mol. Biol. 2016, 88, 1058–1070.

63. Péret, B.; Li, G.; Zhao, J.; Band, L.R.; Voß, U.; Postaire, O.; Luu, D.-T.; Da Ines, O.; Casimiro, I.; Lucas, M.; et al. Auxin regulates aquaporin function to facilitate lateral root emergence. Nat. Cell Biol. 2012, 14, 991–998.

64. Tsugawa, S.; Hervieux, N.; Kierzkowski, D.; Routier-Kierzkowska, A.-L.; Sapala, A.; Hamant, O.; Smith, R.S.; Roeder, A.H.K.; Boudaoud, A.; Li, C.-B. Clones of cells switch from reduction to enhancement of size variability in Arabidopsis sepals. Development 2017, 144, 4398–4405.

65. Sapala, A.; Runions, A.; Routier-Kierzkowska, A.-L.; Das Gupta, M.; Hong, L.; Hofhuis, H.; Verger, S.; Mosca, G.; Li, C.-B.; Hay, A.; et al. Why plants make puzzle cells, and how their shape emerges. eLife 2018, 7.

66. LeGoff, L.; Rouault, H.; Lecuit, T. A global pattern of mechanical stress polarizes cell divisions and cell shape in the growing Drosophila wing disc. Development 2013, 140, 4051–4059.

67. Campinho, P.; Behrndt, M.; Ranft, J.; Risler, T.; Minc, N.; Heisenberg, C.-P. Tension-oriented cell divisions limit anisotropic tissue tension in epithelial spreading during zebrafish epiboly. Nat. Cell Biol. 2013, 15, 1405–1414.

68. Mao, Y.; Tournier, A.L.; Hoppe, A.; Kester, L.; Thompson, B.J.; Tapon, N. Differential proliferation rates generate patterns of mechanical tension that orient tissue growth. EMBO J. 2013, 32, 2790–2803.

69. Louveaux, M.; Julien, J.-D.; Mirabet, V.; Boudaoud, A.; Hamant, O. Cell division plane orientation based on tensile stress in Arabidopsis thaliana. Proc. Natl. Acad. Sci. 2016, 113, E4294–E4303.

70. Gibson, W.T.; Gibson, M.C. Chapter 4 Cell Topology, Geometry, and Morphogenesis in Proliferating Epithelia. In Current Topics in Developmental Biology; Current Topics in Developmental Biology; Academic Press, 2009; Vol. 89, pp. 87–114.

71. Long, Y.; Boudaoud, A. Emergence of robust patterns from local rules during plant development. Curr. Opin. Plant Biol. 2018, 47, 127–137.

72. Vargas-Garcia, C.A.; Ghusinga, K.R.; Singh, A. Cell size control and gene expression homeostasis in single-cells. Curr. Opin. Syst. Biol. 2018, 8, 109–116.

73. Cutler, S.R.; Ehrhardt, D.W.; Griffitts, J.S.; Somerville, C.R. Random GFP::cDNA fusions enable visualization of subcellular structures in cells of Arabidopsis at a high frequency. Proc. Natl. Acad. Sci. 2000, 97, 3718–3723.

74. Stanislas, T.; Hamant, O.; Traas, J. Chapter 11 - In-vivo analysis of morphogenesis in plants. In Methods in Cell Biology; Lecuit, T., Ed.; Cell Polarity and Morphogenesis; Academic Press, 2017; Vol. 139, pp. 203–223.

75. Reuille, P.B. de; Bohn-Courseau, I.; Godin, C.; Traas, J. A protocol to analyse cellular dynamics during plant development. Plant J. 2005, 44, 1045–1053.

76. Legland, D.; Arganda-Carreras, I.; Andrey, P. MorphoLibJ: integrated library and plugins for mathematical morphology with ImageJ. Bioinformatics 2016, 32, 3532–3534.

77. Mirabet, V.; Dubrulle, N.; Rambaud, L.; Beauzamy, L.; Dumond, M.; Long, Y.; Milani, P.; Boudaoud, A. NanoIndentation, an ImageJ Plugin for the Quantification of Cell Mechanics. Methods Mol. Biol. in press.

78. Sassi, M.; Ali, O.; Boudon, F.; Cloarec, G.; Abad, U.; Cellier, C.; Chen, X.; Gilles, B.; Milani, P.; Friml, J.; et al. An auxin-mediated shift toward growth isotropy promotes organ formation at the shoot meristem in Arabidopsis. Curr. Biol. CB 2014, 24, 2335–2342.

79. Kremer, A.; Lippens, S.; Bartunkova, S.; Asselbergh, B.; Blanpain, C.; Fendrych, M.; Goossens, A.; Holt, M.; Janssens, S.; Krols, M.; et al. Developing 3D SEM in a broad biological context. J. Microsc. 2015, 259, 80–96.

80. Kiss, A.; Moreau, T.; Mirabet, V.; Calugaru, C.I.; Boudaoud, A.; Das, P. Segmentation of 3D images of plant tissues at multiple scales using the level set method. Plant Methods 2017, 13, 114.

81. Reuille, P.B. de; Routier-Kierzkowska, A.-L.; Kierzkowski, D.; Bassel, G.W.; Schüpbach, T.; Tauriello, G.; Bajpai, N.; Strauss, S.; Weber, A.; Kiss, A.; et al. MorphoGraphX: A platform for quantifying morphogenesis in 4D. eLife 2015, 4, e05864.

82. Schmidt, T.; Pasternak, T.; Liu, K.; Blein, T.; Aubry-Hivet, D.; Dovzhenko, A.; Duerr, J.; Teale, W.; Ditengou, F.A.; Burkhardt, H.; et al. The iRoCS Toolbox – 3D analysis of the plant root apical meristem at cellular resolution. Plant J. 2014, 77, 806–814.

